# Anatomy promotes neutral coexistence of strains in the human skin microbiome

**DOI:** 10.1101/2021.05.12.443817

**Authors:** Arolyn Conwill, Anne C. Kuan, Ravalika Damerla, Alexandra J. Poret, Jacob S. Baker, A. Delphine Tripp, Eric J. Alm, Tami D. Lieberman

## Abstract

What enables strains of the same species to coexist in a microbiome? Here, we investigate if host anatomy can explain strain co-residence of *Cutibacterium acnes*, the most abundant species on human skin. We reconstruct on-person evolution and migration using 947 *C. acnes* colony genomes acquired from 16 subjects, including from individual skin pores, and find that pores maintain diversity by limiting competition. Although strains with substantial fitness differences coexist within centimeter-scale regions, each pore is dominated by a single strain. Moreover, colonies from a pore typically have identical genomes. An absence of adaptive signatures suggests a genotype-independent source of low within-pore diversity. We therefore propose that pore anatomy imposes random single-cell bottlenecks during migration into pores and subsequently blocks new migrants; the resulting population fragmentation reduces competition and promotes coexistence. Our findings imply that therapeutic interventions involving pore-dwelling species should focus on removing resident populations over optimizing probiotic fitness.

## INTRODUCTION

All host-associated microbiomes live in environments with spatially structured environmental variation generated by host anatomy and physiology. Spatial structure can be considered at multiple length scales—from location along the gastrointestinal tract down to the level of individual crypts and from distant regions on the skin down to the level of individual pores. Revealing the spatial structure of microbial communities is critical for interpreting the coexistence of diverse microbes (Chung et al., 2017; Welch et al., 2016), modeling community assembly and stability (Kerr et al., 2002; Ladau and Eloe-Fadrosh, 2019; Tropini et al., 2017), and predicting the response of microbiomes to therapeutics (Ferreiro et al., 2018; Koskella et al., 2017).

To date, microbiome biogeography studies have largely focused on taxonomic characterization at the species level or higher (Flowers and Grice, 2020; Grice and Segre, 2011; Oh et al., 2016), and intraspecies diversity has received little attention (Rossum et al., 2020; Zhou et al., 2020). Intraspecies diversity can emerge from both the migration of multiple strains to a host and the mutation of individual strains on the host. Sustained diversity arising from both processes has been observed within human microbiomes (Poyet et al., 2019; Zhao et al., 2019; Zhou et al., 2020).

Understanding the forces that generate and maintain intraspecies diversity at both of these levels is critical for the design of precision microbial therapeutics. For example, if adaptive forces like niche partitioning are critical to strain coexistence, then fine-scale manipulation of microbiomes will require understanding the genetic basis of strain success; however, if neutral forces (e.g. priority effects) determine strain composition (Koskella et al., 2017), then therapeutic approaches might depend instead on removal of extant strains.

Evolutionary reconstruction at the whole-genome level, when combined with fine-scaled sampling, provides an opportunity to reveal migration dynamics across a host and the forces maintaining intraspecies diversity (Chung et al., 2017; Jorth et al., 2015; Lieberman et al., 2016). While metagenomic sequencing provides a powerful approach for surveying microbiomes, it does not provide the resolution required for such evolutionary inference. Metagenomic approaches cannot distinguish whether a detected polymorphism reflects recent on-person mutation or the presence of homologous regions among co-colonizing strains. Moreover, metagenomics cannot determine whether a pair of *de novo* mutations occurred on the same or different genetic backgrounds (e.g. 2 mutations each at 20% frequency in the population). While single-cell sequencing can in theory provide information about genomic linkage, current technologies cover only a fraction of the genome and have high error rates. In contrast, culture-based approaches that profile bacterial colonies, each formed from a single cell in the original sample, enable true single-genotype resolution. We therefore use culture-based sequences to obtain the resolution needed for evolutionary reconstruction across an individual host.

The skin microbiome provides an excellent opportunity for studying how spatial structure shapes on-person diversity of commensal microbes (Byrd et al., 2018; Flowers and Grice, 2020) due to the ease of acquiring samples across body sites and its tractability at multiple spatial scales. Here, we focus on *Cutibacterium acnes*, the dominant commensal of sebaceous skin (oily skin of the face and back), because: (1) it is prevalent and abundant across all healthy people; (2) multiple strains of this species stably coexist on each person (Oh et al., 2016); and (3) it can be sampled at multiple spatial scales. *C. acnes* is present on all healthy adults, comprising on average 92% of the bacterial community on sebaceous skin (computed from (Oh et al., 2014)). Despite its name, the role of *C. acnes* in acne vulgaris remains unclear (Dréno et al., 2018; Lomholt et al., 2017; McLaughlin et al., 2019; O’Neill and Gallo, 2018).

Each adult has a unique mix of *C. acnes* strains (Oh et al., 2014). *C. acnes* cells grow faster in anaerobic conditions (Cove et al., 1983) and are thought to consume sebum (the oily substance produced by sebaceous glands at the bottoms of pores) (Brüggemann et al., 2004; Miskin et al., 1997), making the follicles of pilosebaceous units (skin pores) their ideal environment (Fitz-Gibbon et al., 2013; Hall et al., 2018; Leeming et al., 1984). *C. acnes’* residence in anatomical locations that differ greatly in oxygen concentration, nutrient availability, and exposure to the environment raises the possibility that strains display niche specificity. However, it is not yet clear if niche specialization contributes to strain coexistence or why these person-specific populations are resilient to invasion, particularly given the skin’s exposure to the environment.

Here, we report that the anatomy and physiology of human skin promotes substantial intraspecies diversity by segregating the *C. acnes* population across disconnected pores. Strikingly, we find that each skin pore is dominated by a single *C. acnes* strain, despite coexistence of multiple strains within the immediate vicinity. Reduced diversity persists down to the single-nucleotide variant (SNV) level, and phylogenetic reconstruction suggests the presence of single-cell bottlenecks within pores. These bottlenecks cannot be explained by adaptive sweeps, as neutral evolution dominates signatures of on-person evolution. We therefore propose a model in which pore anatomy and physiology gives rise to severe and genotype-agnostic population bottlenecking in skin pores, thereby reducing interstrain competition and promoting the maintenance of intraspecies diversity via non-selective means. More broadly, these findings present a framework for using SNV-level spatial biogeography to uncover migration dynamics at the subspecies level and highlight the capacity of anatomy to shape the ecology and evolution of commensal microbes.

## RESULTS

### *C. acnes* biogeography at unprecedented spatial and genetic resolution

In order to capture the biogeography of *C. acnes* on sebaceous skin of healthy people, we collected samples across multiple length scales (Figure 1). At the finest scale, we collected material from inside single sebaceous follicles—where most *C. acnes* growth is thought to take place–using comedone extractors and pore strips (pore samples; Methods). We incidentally collected samples that included material from multiple adjacent follicles (multipore samples; Methods). In addition, we collected samples on a coarser spatial scale (forehead, nose, left/right cheek, chin, shoulder, back quadrants) by scraping a long toothpick back and forth over a large sebaceous skin region (scrape samples). In total, we collected 300 samples from 16 healthy adults, including 145 pore samples from 5 of these subjects (Tables S1-S2).

**FIGURE 1:**
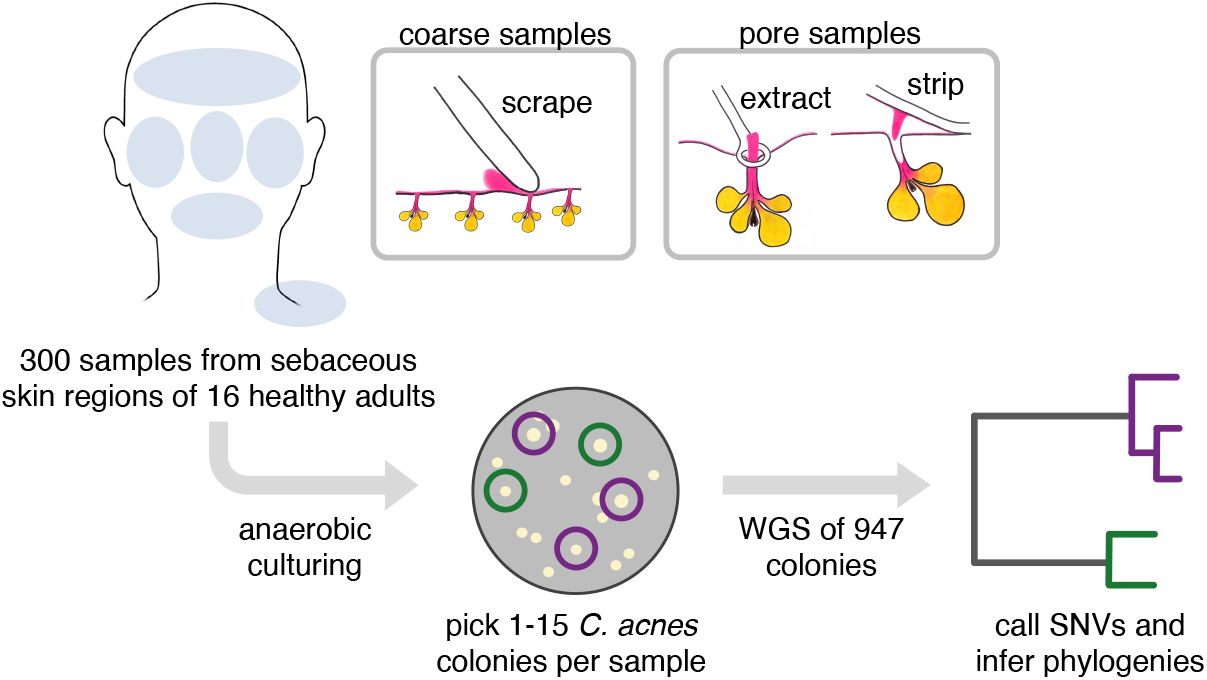
*Cutibacterium acnes* biogeography at high spatial resolution and high genetic resolution. We collected *C. acnes* colonies from sebaceous skin regions (forehead, nose, right cheek, left cheek, chin, shoulder, back) from 16 healthy adult subjects. Samples were acquired at coarse spatial resolution with a toothpick (scrapes; N=155) and fine spatial resolution (pore extracts and pore strips; N=145). This approach enabled us to examine *C. acnes* biogeography with spatial resolution down to a single pore (sebaceous follicle). To understand how these samples were related to each other, we performed whole genome sequencing on 947 colonies (1-15 per sample), each of which represents the genetic content of a single cell that originated on the skin of one of our subjects. This approach enabled us to examine *C. acnes* biogeography with genetic resolution down to a single nucleotide variant (SNV).

Immediately after sampling, we streaked the collected material onto solid media and incubated in an anaerobic environment that favors *C. acnes* growth (Methods). We randomly selected 1-15 colonies per sample with colony morphology consistent with *C. acnes* for whole genome sequencing (Figure 1). All together, we obtained 947 high-quality genomes that passed purity and coverage filters (Methods).

### *C. acnes* communities on individuals arise from multiple colonization events

We first classified our colonies using an established typing scheme (Scholz et al., 2014) (Methods); the 7 strain types represented in our dataset cover the majority of known *C. acnes* diversity. Consistent with previous work (Lomholt et al., 2017; Oh et al., 2014), we find that multiple *C. acnes* strain types typically reside on an individual person. However, the phylogenetic distribution of strain types present varies considerably from person to person (Figure 2A).

**FIGURE 2:**
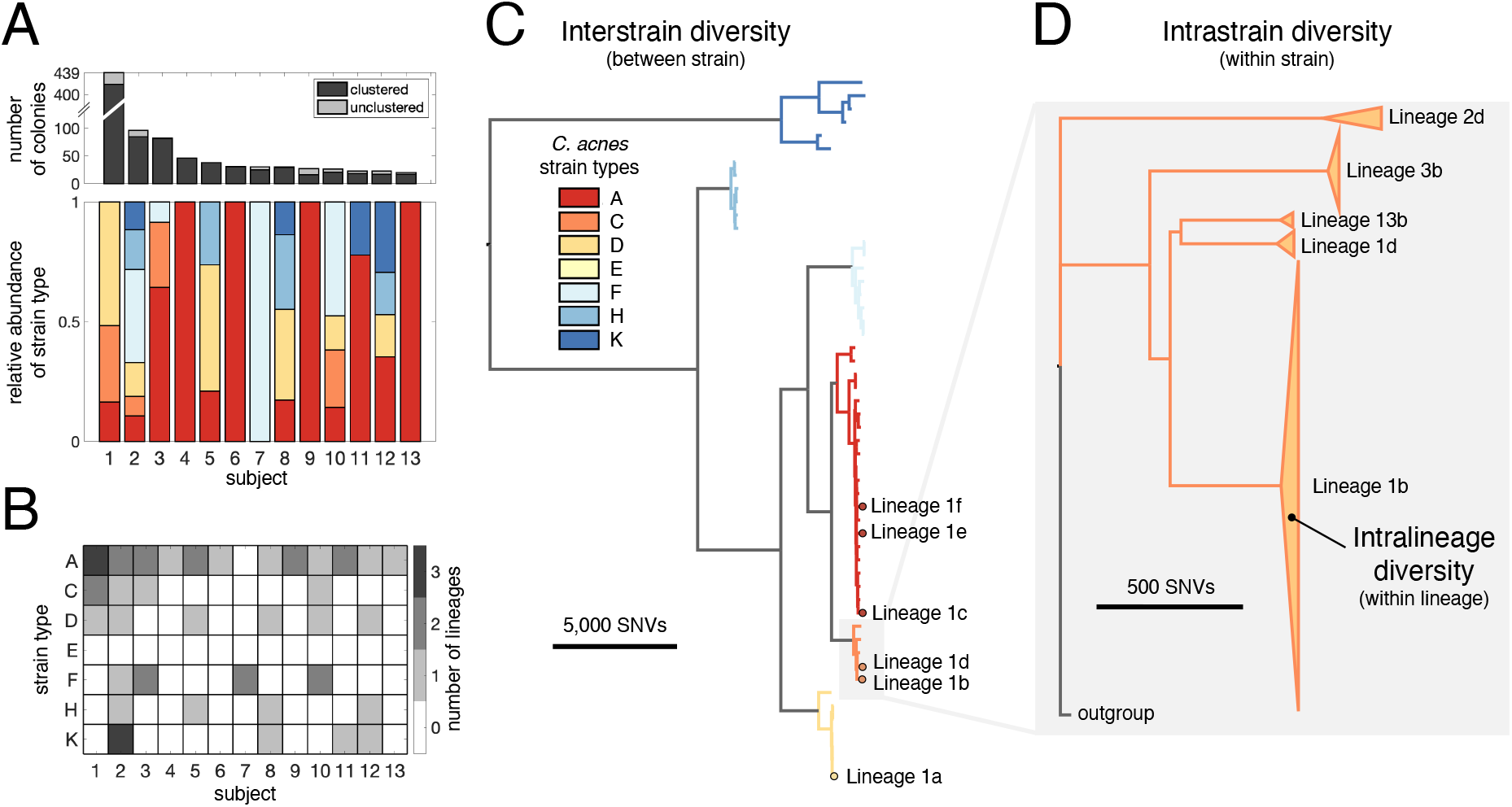
*C. acnes* lineages from distinct colonization sources coexist on individuals. (A) Multiple coexisting strain types of *C. acnes* typically reside on the skin of healthy adults, with the strain-level composition varying between individuals (lower panel). The number of *C. acnes* colonies per subject is shown in the top panel, with the number of unclustered colonies in light gray. Only subjects for whom at least 20 colonies passed quality filters are shown (Methods). The relative abundances of *C. acnes* strain types on each subject is based on an established *C. acnes* strain typing scheme (Scholz et al., 2014). (B) Subjects often have multiple lineages belonging to the same strain type. We note that the number of lineages detected on an individual is sensitive to sampling depth (Figure S1). (C) Phylogenetic relationship of 53 *C. acnes* lineages detected across all subjects is shown and colored by *C. acnes* strain type. All six distinct coexisting lineages found on Subject 1 are highlighted. (D) A zoom-in of strain type C illustrates that lineages within a strain type are separated by large genetic distances relative to intralineage diversity. The heights of triangles are proportional to the number of colonies in each lineage and their widths represent the extent of intralineage genetic divergence. Lineages are named by subject number and then indexed by size within each subject using lowercase letters.

To assess whether colonies of the same strain type might originate from independent colonization events, we quantified genomic divergence using a reference-based approach and focused primarily on single nucleotide variants (SNVs) (Methods; Figure 1). This approach captures the vast majority of the intraspecies variation because *C. acnes* has a small accessory genome (~10% variation between strain types), primarily composed of genomic islands that do not vary among colonies of the same strain type (Brzuszkiewicz et al., 2011; Scholz et al., 2016; Tomida et al., 2013) (Table S3). Plasmids make up most of the mobile gene content, and 31% colonies have evidence of a plasmid (Brüggemann et al., 2012; Kasimatis et al., 2013) (Methods; Table S2).

We found that colonies from the same individual, but not different individuals, were often very closely related: while >99% of pairs of colonies from different subjects were separated by >35 SNVs across their genomes, 28% of pairs of colonies from the same subject had genetic distances below this threshold (Figure S1). This disparity suggests that closely related colonies emerge from on-person diversification from a recent ancestor on that individual (Zhao et al., 2019; Zhou et al., 2020). We therefore clustered colonies into lineages based on genetic distances, resulting in 54 lineages — each of which contains colonies from only one subject (Methods; Figure 2B–D; Table S4). Because we imposed a minimum cluster size of 3 colonies, some colonies do not belong to any cluster; these represent either low-abundance genotypes or transient non-resident genotypes from the external environment. While colonies within a cluster have diverged only 0-26 SNVs from their lineage’s inferred common ancestors (median within a cluster; Table S5), lineages are much more divergent from one another (Figure 2C–D).

The clustering of colonies into lineages allowed us to estimate the number of colonization events on each individual. Each lineage might represent a distinct colonization event (Zhao et al., 2019), or a lineage might reflect multiple colonization events if a person is colonized by multiple closely related genotypes (e.g. multiple genotypes from a parental lineage transferred to a child). Therefore, the number of *C. acnes* lineages found on a person represents the minimum number of *C. acnes* genotypes that successfully colonized a person. We note that we underestimate lineage coexistence on many subjects, as most were not sampled exhaustively (Figure S1, Table S1). Intriguingly, we often detected multiple lineages of the same strain type on an individual subject (Figure 2B), demonstrating that an individual host can be colonized by the same strain type multiple times. In the most extreme case, Subject 2 has been colonized at least 9 distinct lineages from 6 strain types.

### Coexistence of *C. acnes* strain types does not arise from specificity to anatomical niches

To test if strain types coexist because they are equally competitive, we measured their growth rates *in vitro.* Even in the simplest of laboratory conditions, we noticed substantial differences between strains originating from the same person. We assessed growth rates for 18 colonies from the 3 most abundant lineages on Subject 1 (the most intensively sampled subject), all from different strain types and cultured from the same timepoint (Figure S2). We find that growth rates vary up to 30% across colonies (p < 10^−12^, ANOVA), with variation apparent both within and across lineages. We therefore sought to identify what enables *C. acnes* strain types with different competitive abilities to coexist *in vivo.*

The stable coexistence of diverse *C. acnes* strain types might arise from niche specialization to anatomical features. In particular, the environment on the skin surface differs dramatically from that inside skin pores in terms of oxygen concentration, nutrient availability, and other factors (Adamson and Lipoff, 2021; Plewig et al., 2019). We therefore looked for differences in strain types when sampling directly from the follicle of a pore (extract and pore strip samples) as compared with sampling across the skin surface (scrape samples). However, we did not observe strain specificity to the skin surface vs skin pores on Subject 1 (Figure 3A) or across subjects (Figure 3B). This suggests that *C. acnes* strain types are not specialized to either the pore or the skin surface environment and that diverse strain types are similarly competitive across anatomical settings.

**FIGURE 3:**
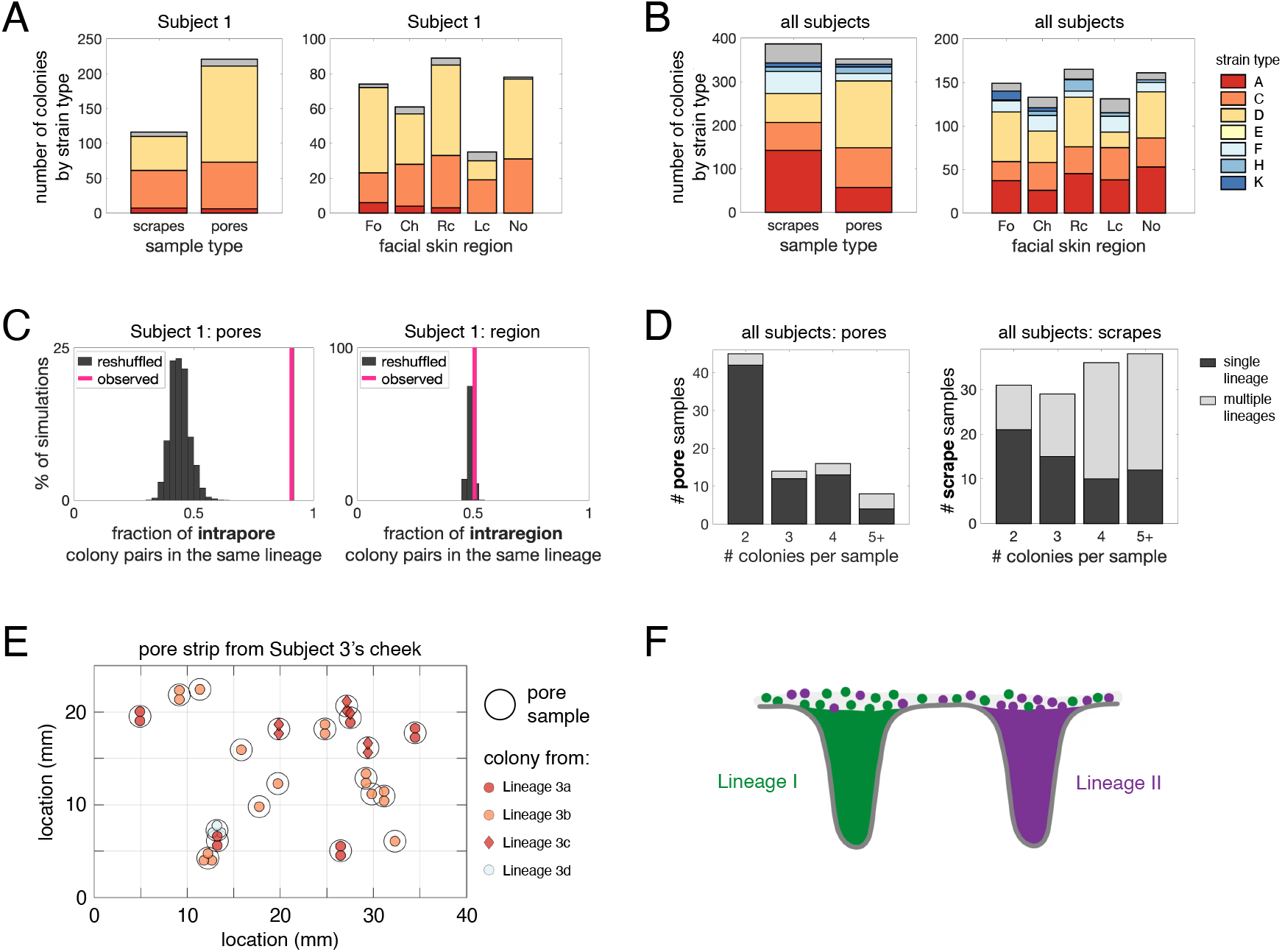
*C. acnes* lineages are spatially segregated into different pores despite coexistence on skin regions. (A-B) Niche specificity does not explain coexistence of strain types on an individual. Strain types follow the same color scheme as in Figure 2. (A) *C. acnes* strain types on Subject 1 are not specific to sample type (coarse scrape samples vs pore samples) or to skin regions (Fo=forehead, Ch=chin, Rc=right cheek, Lc=left cheek, No=nose). (C) Multiple strain types or lineages do not typically coexist within the same pore. Pairs of colonies from the same pore belong to the same lineage 91% of the time (pink line). In contrast, randomly reshuffled colonies from the same pore belong to the same lineage only 44% of the time (gray histogram; p<0.0001). In comparison, colonies originating from different pores within the same facial region are only slightly more likely to be from the same lineage than when compared to a random model (p=0.03). (D) The enrichment of single lineages within individual pores persists across all subjects. Colonies from the same pore sample typically belong to the same lineage (left), whereas colonies from the same coarse samples often belong to more than one lineage (right). Analyses in (C) and (D) exclude pore samples originating from more than one follicle. (E) A pore strip from Subject 3 (left cheek section) illustrates how pores housing different lineages can be in close vicinity to each other. Each black circle represents a single pore sample, and each interior symbol represents a colony from that follicle. Symbol colors indicate strain type and symbol shapes indicate lineage. (F) These findings demonstrate that lineages are spatially segregated into different pores, despite lineage coexistence within skin regions.

We next explored the possibility that some strain types are better adapted to particular skin regions (e.g. nose vs forehead), but we find no signature of specialization on the face. Diverse strain types coexist in close proximity within facial skin regions on Subject 1 (Figure 3A). This pattern holds across subjects, where we do not find any enrichment patterns indicating facial region specificity (Figure 3B). This lack of specialization on the face is consistent with previous metagenomic and culture-based studies (Lomholt et al., 2017; Oh et al., 2014, 2016). Some subjects, however, harbor substantial compositional differences in their *C. acnes* strain types between the face and back, a pattern also apparent in publicly available metagenomic data (Figure S3) (Oh et al., 2014). Interestingly, we find no consistency in which strain types are enriched on faces and backs. This lack of consistency argues against a ‘back-adapted’ or ‘face-adapted’ strain and instead implicates neutral forces such as limited migration or priority effects (forces that favor early colonizers over new migrants). Together, these findings support a model in which *C. acnes* strain types are not specialized to specific anatomical regions.

### Each skin pore is dominated by only one lineage

The lack of niche specialization to anatomical features raises the question of how the skin environment prevents strain types from outcompeting each other. We next investigated fine-scale spatial resolution, focusing on the lineage level and on samples obtained from pore follicles (pore strips and extracts).

At the level of individual pores, we observe a striking absence of diversity (Figures 3C–D). This can be most clearly seen by close examination of Subject 1, from whom we sampled the greatest number of pores. Pairs of colonies from Subject 1 originating from a pore belong to the same lineage at a significantly higher rate than would be expected if genotypes were randomly distributed (91% vs 44%, p<.0001; Figure 3C). Notably, from the most densely sampled pore, 11 of 11 colonies are from the same lineage (sample 5 in Lineage 1a). Across all subjects, we observe similar trends of lineage-enrichment within pores, despite lineage coexistence within scrapes (Figure 3D). This segregation persists even when pores are closely spaced; for instance, in a 1 square cm section of a pore strip from Subject 3, we found 3 different lineages, despite each pore containing only a single lineage (Figure 3E).

Although low within-pore diversity (Figure 3F) might arise from sampling methods that only capture representatives from a part of the follicle, we note that previous work using light microscopy to image skin biopsies after blackhead extraction suggests that extractions are capable of removing the majority of the follicular contents (Plewig, 1974).

### Monocolonization of pores results from neutral bottlenecks

Spatial segregation of *C. acnes* lineages in skin pores could arise from priority effects or from pore-specific selection shaped by the host or other microbes. We reasoned that these mechanisms would result in different degrees of within-pore diversity when examined at the whole-genome level, as well as different signals of adaptive evolution. Exclusion via priority effect or adaptive sweep within a pore would result in a single genotype within each pore, while selection for members of a particular lineage would sometimes result in coexistence of distinct migrants of the same lineage.

At the level of individual SNVs (Methods; Table S5), we find a striking lack of *C. acnes* intrapore diversity, with colonies from the same pore clustering tightly together on the phylogeny (Figure 4; Figures S4-6). Colonies from the same pore often form monophyletic clades, and in some cases share mutations not detected anywhere else or rare plasmid variants (Figure S7). Moreover, metrics of intrapore diversity are extremely low relative to each lineage’s total diversity, as assessed by genetic distances to various inferred most recent common ancestors (MRCA). Colonies in Lineage 1a (the largest lineage from Subject 1) from single pore samples have on average less than 1 mutation since their intrapore MRCA, whereas pairs of pores from this lineage typically have 4-8.5 mutations (25%-75% percentiles) since their interpore MRCA (Figure 5A). This pattern of extremely low intrapore diversity, in both absolute and relative scales, is consistent across lineages and subjects (Figure 5A; Figures S4-6; Table S5).

**FIGURE 4:**
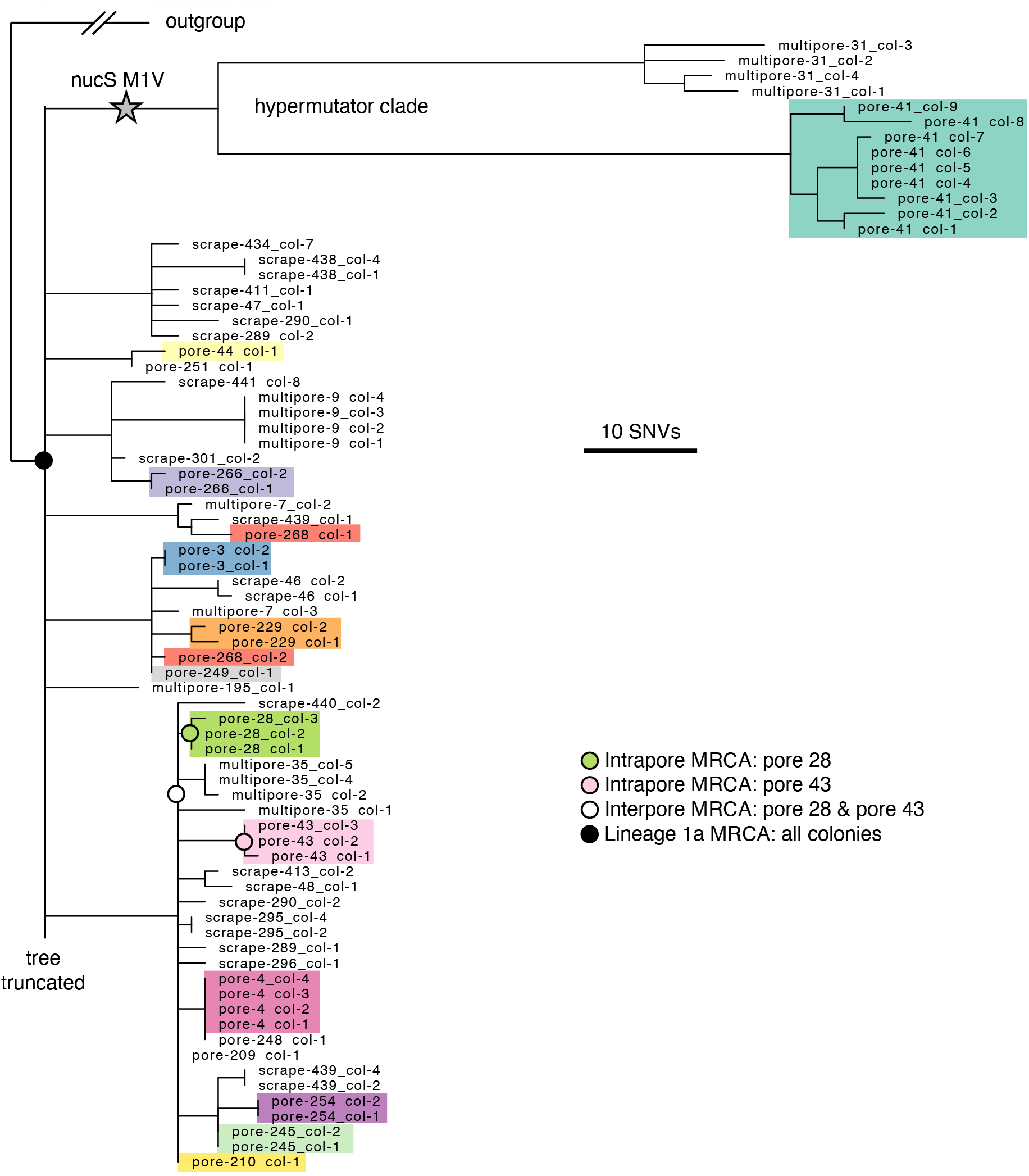
Each pore harbors only a small fraction of intralineage diversity. Maximum parsimony tree of the most abundant lineage on Subject 1, Lineage 1a, in which each leaf represents a single colony. Colonies are colored by pore (excluding multipore samples and pore samples with only one colony), emphasizing low within-pore diversity. The long branches at the top of the phylogeny display a hypermutator phenotype (Figure S4). For any given non-hypermutator pore, the mean genetic distance of colonies to the pore’s inferred most recent common ancestor (MRCA) is usually less than 1 SNVs (median across pores: 0 SNVs; 25%-75% percentiles: 0 to 1.1 SNVs). Four example inferred ancestral genotypes are marked on the tree. Due to space limitations, the tree is truncated; see Figure S4 for the complete tree.

**FIGURE 5:**
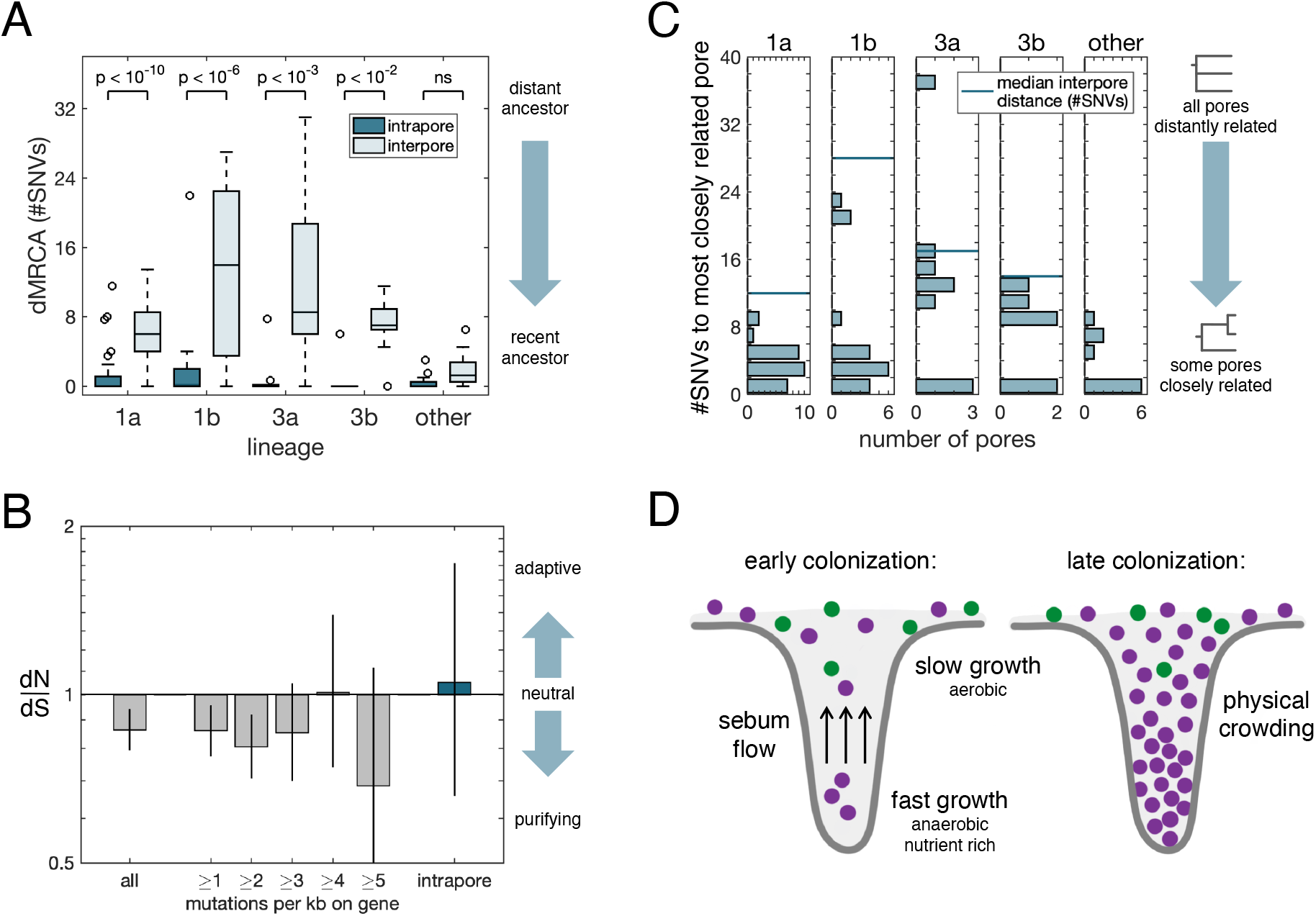
Neutral forces give rise to population bottlenecks during pore colonization. The pattern of small intrapore dMRCAs (distance to the MRCA, averaged across colonies) as compared with interpore dMRCAs (average distance from the two pore MRCAs to their interpore MRCA) is consistent across subjects and lineages (Wilcoxon rank-sum test with Bonferroni correction). This indicates that pore populations are subject to recent strong bottlenecks and are ecologically isolated from each other. Analysis for (A) and (C) included all single pore samples (excluding hypermutators in Lineage 1a); pores from lineages containing fewer than five such samples are grouped as “other”. (B) Across all within-lineage SNVs, dN/dS (the ratio of nonsynonymous to synonymous mutations, relative to a neutral model; Methods) is slightly negative, indicative of purifying selection. Values of dN/dS for genes with high mutational densities among within-lineage mutations are consistent with a neutral model, as is dN/dS for mutations inferred to have occurred inside pores. These findings suggest that neutral evolution dominates *C. acnes* evolution on individuals and that adaptive sweeps are not responsible for low within-pore diversity. Error bars indicate 95% CIs. See Figure S10 for a more detailed analysis. (C) Pairs of pores on a person often share very recent common ancestry, suggesting that neutral bottlenecking occurred during a recent pore colonization or recolonization event. The genetic distance between two pores is equal to twice the interpore dMRCA. Given that the number of pores sampled per subject was vastly smaller than the total number of pores on a person, these values underestimate the commonality of shared mutations between pore populations. (D) Pore physiology may contribute to the neutral bottlenecking process (see also Figure S11). Sebum flow out of the pore, the environmental gradient along the length of the pore, and physical crowding all make it more difficult for a cell to colonize a pore.

Although the molecular clock rate for *C. acnes* is not known and we were unable to accurately measure it (Figure S8), all reported bacterial molecular clocks from human infection or colonization range between 0.5 SNVs/genome/year and 30 SNVs/genome/year (Didelot et al., 2016; Zhao et al., 2019). Therefore, our observation of low intrapore diversity (median 0 SNVs since pore MRCA, 25%-75% percentiles: 0-0.6 SNVs; Methods) suggests that the population within each pore typically descended from a single cell about 1 year ago and hints that priority effects may be important to the exclusion of other strain types.

There are two pore samples in Lineage 1a that have diverged further from the lineage MRCA (45 and 56 SNVs vs a mean of 9 SNVs; Grubbs outlier test) and harbor more intrapore diversity. We suspected that this excess diversity might be due to hypermutation, an accelerated mutation rate that is common in laboratory experiments (Sniegowski et al., 1997) and *in vivo* (LeClerc et al., 1996), usually caused by a defect in DNA repair (Oliver, 2010). Consistent with this hypothesis, these colonies share a mutation that eliminates the start codon of the *nucS* gene, which encodes for an endonuclease critical for the repair of transition mutations (Castañeda-García et al., 2020; Ishino et al., 2018). Indeed, we observe an enhancement in the ratio of transition to transversion mutations in the hypermutator clade (Figure S4). This finding suggests that these pores were physiologically similar to other pores, and that an increased mutation rate enabled *C. acnes* to accumulate more diversity between the most recent single-cell ancestor and sampling.

The finding of a recent single-cell ancestor for each pore is particularly surprising given that single pores contain on average 50,000 colony-forming units of *C. acnes* (max 10^8^ CFU; Figure S9) (Claesen et al., 2020). Such large population sizes generally limit the speed of neutral genetic drift (Hartl and Clark, 2006); classic models of neutral evolution predict that it would take over 100,000 bacterial generations (in this case, likely hundreds of years) for a neutral mutation to sweep a population of this size. Therefore, our observations suggest the presence of either conditions that enhance genetic drift or adaptive mutational sweeps that swiftly purge diversity.

To test if adaptive sweeps might be responsible for purging diversity inside pores, we examined within-lineage mutations for evidence of past adaptation. Parallel evolution is a common signature of adaptation in bacteria that manifests as an enrichment of mutations in genes or pathways under selection relative to a neutral model (Lieberman et al., 2011; Zhao et al., 2019). However, we detected no cases of parallel evolution across all 2,445 *de novo* mutations in coding regions, across mutations occurring on a subject, across mutations occurring within a lineage, or among intrapore mutations (Figure S10, Methods). Moreover, we identified a depletion of nonsynonymous (amino-acid changing) mutations relative to a neutral model among all *de novo* mutations (dN/dS < 1, Figure 5B), which is invariant to the number of times a gene was mutated, the inferred age of a mutation, or functional pathways (Figure S10). These signals suggest that neutral forces shape on-person evolution of *C. acnes* and argue against selective sweeps as the driver of within-pore bottlenecks. Instead, we propose that low within-pore genetic diversity stems from frequent, neutral population bottlenecks induced by pore anatomy and physiology.

### Pore anatomy and physiology are sufficient to create bottlenecks during colonization

We next asked if these recent population bottlenecks occurred long after pore populations were established, or, instead, during recent migration into a pore. If pore populations are segregated for long periods of time, the recent bottlenecks observed here would reflect only the most recent bottleneck in a series of in-pore bottlenecks; in this case, sequential intrapore sweeps would create large genetic distances between the MRCAs of each pore. Instead, we find that most pores have a closely related population in another pore, with many pairs of pores sharing SNVs inferred to have occurred recently (Figure 5C; Figures S4-6). These findings are consistent with recent transmission of genotypes between pores. Combined with our observations of young populations within pores (Figure 5A), the finding of recent common ancestors between pores supports a model in which neutral bottlenecking occurred during recent pore colonization or recolonization events.

We propose that pore physiology can create such bottlenecks (Figure 5D; Figure S11). We modeled the process of pore colonization, using published values for relevant physiological parameters (Butcher and Coonin, 1949; Cove et al., 1983; Plewig, 1974) and the assumption that most *C. acnes* growth occurs in the favorable conditions at the bottom of pores. First, since *C. acnes* is not motile (Brüggemann et al., 2004), it must rely on growth and diffusion in order to reach the bottom of a pore. Estimations of the diffusion coefficient of a bacterial cell in sebum and of the sebum flow speed suggest that most potential colonizers are quickly pushed out of the follicle by the sebum flow (Butcher and Coonin, 1949; Plewig, 1974); it is rare for a cell to remain in a pore for more than one doubling-time. Second, *C. acnes* cells likely cannot proliferate rapidly until they reach lower depths in the pore, where the environment is anaerobic and nutrient rich due to sebum production (Cove et al., 1983; Flowers and Grice, 2020). Third, solid obstacles, including bacterial mass and dead human cells (Jahns and Alexeyev, 2014; Plewig et al., 2019), embedded in sebum will further slow diffusion, making it even more difficult for potential invaders to colonize. In this way, pore physiology could enable a lucky single cell to found a pore’s resident population, with abundant growth at the bottom of the pore blocking new migrants.

Despite small distances between some pore MRCAs, the MRCA for each lineage as a whole is substantially older (Figure 5C). These data are consistent with a model in which pore populations studied here were established long after a given lineage initially migrated onto a subject’s skin. We therefore propose that these colonization events may represent pore recolonization events following a disturbance to the underlying community, perhaps caused by the immune system, phage predation, or physical clearing.

### Pores are colonized by *C. acnes* genotypes from distant locations

To understand migration dynamics across pores, we turned to pore strip data, where each pore sampled has defined spatial coordinates (Figure S12; Table S6). In the case that pores are colonized preferentially by their neighbors, we would expect to see spatial confinement of genetic variants that emerged recently. However, similar to our previous observation that lineages themselves are not specific to certain facial skin regions, we find that closely related pores can be separated by large physical distances (e.g., Figure S5). To assess this quantitatively, we created a neutral model in which spatial coordinates are randomly shuffled and assessed whether pores with closely related genotypes were more likely to be in the vicinity of each other than by random chance, and we find no evidence of spatial confinement at the SNV level (Figure S13). This finding suggests that the timescale for a new genotype spreading across facial skin regions is faster than the timescale for further genetic diversification. This is consistent with a model in which *C. acnes* cells primarily grow within pores and are transferred across the skin to newly opened pores via long-range dispersal mechanisms (e.g. washing or touching).

### Skin pores promote coexistence and stability of extant *C. acnes* lineages

Altogether, our results support a model in which bottlenecking in skin pores and, therefore, skin anatomy and physiology, play a major role in *C. acnes* on-person ecology. As a consequence of severe spatial segregation into island-like units, *C. acnes* populations in different pores do not rely on the same resources for growth. Theoretical work has proposed that such spatial segregation promotes neutral coexistence by reducing the strength of ecological interactions (Coyte et al., 2015). We propose that the reduction in competition promoted by isolated pores can help explain the coexistence of *C. acnes* lineages on skin.

Moreover, the priority effects created by pores may help explain the surprising observations that an individual’s strain types are stable over time despite the skin’s exposure to the outside world (Oh et al., 2016). First, the physiology of pores insulates their *C. acnes* populations from the external environment. Moreover, sebum flow ensures that *C. acnes* cells on the skin surface originating from pores outnumber those originating from the environment. Consequently, already established lineages will have a higher likelihood of colonizing a newly available pore. Longer timeseries data will be crucial to understanding the extent to which pores stabilize community dynamics over the host’s lifetime.

Taken together, our findings support a model in which skin pores play a critical role in *C. acnes* ecology. Skin pores provide an environment well-suited for *C. acnes* growth, but population bottlenecking limits the amount of genetic diversity each pore harbors. As a consequence, skin pores both reduce competition between strain types via spatial segregation and favor the existing community via a priority effect. These forces work together to create a stable skin population resilient to invasion.

## DISCUSSION

### Skin pores promote strain coexistence via neutral processes

In this work, we have shown that skin anatomy strongly influences the generation and maintenance of intraspecies diversity in *C. acnes*, a prevalent and prominent commensal on human skin. Our culture-based approach and fine-scaled sampling methods enabled us to examine *C. acnes* biogeography with resolution down to single SNVs and single skin pores (Figures 1–2). This resolution was essential for uncovering that the *C. acnes* population in a single skin pore is extremely bottlenecked (Figures 3–5). We propose that this bottlenecking can explain the stable coexistence of diverse *C. acnes* populations on individual adults (Oh et al., 2016), despite differences in strain fitness and despite the skin’s exposure to the environment.

We did not sample enough individuals in this study to characterize how different skincare regimens or history of treatment for acne might alter *C. acnes* biogeography. As we only studied adult subjects without acne vulgaris, future studies will be needed to understand implications of these findings for acne (Dréno et al., 2018; Lomholt et al., 2017; McLaughlin et al., 2019; O’Neill and Gallo, 2018). However, we note that we found similar patterns across all subjects studied, suggesting that our observation of low within-pore *C. acnes* diversity is not driven by a specific skincare regimen.

Future studies will be needed to understand if the findings we report for *C. acnes* are relevant to other skin commensals, and, more broadly, if crypt-like structures promote intraspecies diversity in other microbiomes. Intriguingly, our dataset includes 3 pore samples from which we cultured multiple clonal *Cutibacterium granulosum* colonies (Figure S14), hinting that the process leading to low within-pore *C. acnes* diversity may also apply to other related pore-dwelling species on human skin (Mak et al., 2013). However, we do not necessarily expect these patterns to hold for *Staphylococcus epidermidis* and related species, which are thought to grow primarily at the tops of pores and on the skin surface (Plewig et al., 2019).

Beyond the skin, the crypts of the large intestine have been shown to promote priority effects among *Bacteroides* in mice (Whitaker et al., 2017). However, at least for *Bacteroides fragilis*, toxin secretion is thought to be integral to exclusion of other strains (Hecht et al., 2016); this non-neutral filtering mechanism may explain why strain co-existence in this species is rare (Garud et al., 2019) despite the presence of crypts and priority effects. We speculate that the importance of crypt-like structures in maintaining intraspecies diversity will depend both on microbial strategies and whether the particular anatomical and physiological conditions induce single-cell bottlenecks.

### Role of skin pores in the balance of neutral and adaptive evolution

Our finding that SNV-driven adaptive evolution is exceedingly rare in *C. acnes* evolution—to the point where it is undetectable here (Figure 5; Figure S10)—is surprising in light of recent reports of rapid adaptive evolution in other stable members of human microbiomes (Poyet et al., 2019; Zhao et al., 2019). While low population sizes can limit adaptive evolution (Ghalayini et al., 2018; Hartl and Clark, 2006), *C. acnes* populations on individuals can reach up to 10^10^ cells, suggesting ample potential for on-person evolution. One possible explanation for our observation is that exceedingly few beneficial mutations remain to be explored (Wielgoss et al., 2013; Wiser et al., 2013). For example, the skin microenvironment might be relatively stable compared to the variable environment of the human gut, selective pressure from bacteria might be limited by the relatively low complexity of the microbial community on skin (Oh et al., 2014), or follicle structure or sebum flow might limit phage predation (Lourenço et al., 2020)—all of which would result in fewer opportunities for adaptation for skin commensals.

Alternatively, it is possible that our observations of largely neutral evolution arise from the dominance of stochastic forces on the skin. To that end, we hypothesize that the physical structure of pores may create an environment in which luck and location—rather than genomically-encoded fitness—predict success, therefore limiting the adaptive potential of *C. acnes* on individual people. Bottlenecking suppresses selective forces by both reducing competition between cells with different genotypes and by introducing randomness in which cells get to proliferate (Barrick and Lenski, 2013; Lieberman et al., 2005; Tenaillon et al., 2016). In addition, genetic drift may be favored because the number of cells that are actually growing might be substantially smaller than the census population (e.g. if bacterial replication were restricted to the very bottom of the follicle) (Hartl and Clark, 2006). In the case of a narrow growth region, physical crowding of cells inside a pore (Jahns and Alexeyev, 2014; Plewig et al., 2019) may exclude beneficial mutants from the growth layer (Schreck et al., 2019). These proposed mechanisms emphasize how host anatomy has the potential to suppress selective forces and tip the balance toward more neutral outcomes. They also raise an interesting question of whether these structures have evolved because limiting commensal evolution is beneficial to the host (Foster et al., 2017).

### Implications for microbial therapeutics

Understanding how host anatomy and physiology influence strain-level composition in microbiomes is critical to the design of precision microbiome therapeutics—particularly those that are intended to engraft into the existing community or remove a member of that community. This study of skin pores exemplifies how host anatomy can contribute to strain-level coexistence and stability via neutral means, with implications for the development of microbiome-based therapeutics (Costello et al., 2009; Paetzold et al., 2019; Schmidt, 2020). In particular, these results suggest that the ability of a probiotic strain to engraft on sebaceous skin will hinge less on the probiotic strain’s competitive fitness and more on efficient removal or destabilization of the existing community prior to treatment.

Here, we have shown that evolutionary reconstruction of mutations—including neutral ones—at the SNV scale reveals migration dynamics in the microbiome and provides insight into the processes by which genetic diversity is maintained. We anticipate that future studies applying similar evolutionary approaches to other microbes will accelerate development of the mechanistic understanding needed for precision microbiome engineering.

## MATERIALS AND METHODS

### Human subjects and sample collection

Healthy adult subjects who had not taken antibiotics in the past 3 months were recruited under a protocol approved by MIT’s Institutional Review Board. Subjects were asked to wash their face with gentle soap prior to sampling to enrich for resident bacteria. In order to sample from diverse anatomical features—including skin pores (sebaceous follicles) and the skin surface— three sampling methods were employed (Figure 1).

Scrape samples were collected using a long sterile toothpick to survey bacteria from both the surface and the tops of pores within a given facial region. Scrape samples were collected from each subject at 8 standardized regions: forehead, left cheek, right cheek, chin, upper right back, upper left back, lower left back, and lower right back. From some subjects, additional scrape samples were collected (Tables S1 and S2). Each toothpick was dragged at an angle using 1-2 inch strokes about 10 times over the region to be sampled, turning occasionally to maximize biomass collection. Each toothpick was then used to immediately inoculate Brucella Blood Agar plates (Hardy Diagnostics) and spread for single colonies using fresh inoculator loops.

For select subjects, samples from inside pore follicles were collected using a comedone extractor or pore strips. Pilosebaceous units (pores) to be sampled via comedone extraction were identified visually as blackheads or whiteheads, and a sterilized comedone extractor was used to apply pressure to the surrounding area of skin. Most extracts removed contents from a single pore as a semi-solid plug. However, some attempts resulted in the extraction of contents from multiple follicles; these samples were labeled as ‘multipore’ samples. For single-pore and multipore samples, a sterile plastic inoculator loop was then used to transfer the pore contents to a Brucella Blood Agar plate. This first inoculator loop was struck multiple times on the plate to disturb the follicular plug, which was then struck out for single colonies as above. When multiple pores were extracted simultaneously, contents from all extracted pores were processed together and the sample was labeled as containing contents from multiple pores. Some extracts (indicated in Table S1) were processed like pore strip samples (below) in order to conduct amplicon sequencing as well.

For pore strips, a commercially available product (Charcoal Blackhead Mask, Mengkou) was applied to the cheeks, nose, and forehead and allowed to dry. The dried film was carefully peeled off and segments were placed into sterile petri dishes for processing. Spatial coordinates for pore strip samples are available in Table S6. Under a dissection microscope, individual extracts were plucked off using sterilized forceps and placed into individual wells of a microplate containing 50 μl of QuickExtract buffer (EpiCentre). Extracts were disturbed by pipetting up and down (samples did not completely dissolve even after mixing). A 5 uL aliquot was used to inoculate a Brucella Blood Agar plate and struck for single colonies. The remainder was used for amplicon sequencing as described below (see 16S amplicon sequencing).

### Culturing and single-colony sequencing

Culture plates were incubated in an anaerobic environment at 37°C for 5-7 days to enrich for *C. acnes*. Random colonies suspected to be *C. acnes* based on colony morphology were selected for further profiling. From most samples, up to 4 colonies were chosen for further processing; additional colonies were chosen on a few samples for further depth (see Table S2 for details on colonies that passed all filters). Selected colonies were resuspended in 200 μL of PBS, and 150 μL of the material was used for gDNA extraction. To obtain more pure freezer stocks, a small subset of colonies was restreaked prior to making freezer stocks; these isolates were used for growth-rate analysis and long-read sequencing and are indicated in Figure S2 and below (see *C. acnes* plasmid analysis), respectively. The remainder of the colony was mixed with glycerol to reach a final concentration of 20% and frozen at −80°C. DNA was extracted in 96-well plates using the PureLink DNA Extraction Kit (Invitrogen), using instructions for gram-positive bacteria, with the exception of longer incubations times (12 hours lysozyme step; 3 hours proteinase K step) and elution into a smaller volume (20 ul). Genomic libraries for Illumina sequencing were prepared according to a previously described protocol (Baym et al., 2015). Libraries were pooled and sequenced on Illumina NextSeq and HiSeq using 75-bp paired end reads to an average depth of 76 reads for colonies passing eventual filters (Table S2).

### Clustering colonies into lineages

Colonies were clustered into lineages using SNV calls from an alignment-based approach with pipelines implemented in snakemake (v5.6.0) (Mölder et al., 2021) and Matlab (v2018a) (code will be available at: https://github.com/arolynconwill/cacnes_biogeo). Adapters were removed using cutadapt (Martin, 2011) and reads were trimmed using sickle (v1.33; -q 20 -l 50 -x -n) (Joshi and Fass, 2011). Next, reads were aligned using bowtie2 (v2.2.6; -X 2000 --no-mixed -- dovetail) against *Cutibacterium acnes* C1 (RefSeq NC_018707) (Langmead et al., 2009; Minegishi et al., 2013). Candidate single nucleotide variants were called using samtools (v1.5) mpileup (−q30 -t SP −d3000), bcftools call (-c), and bcftools view (−v snps −q .75) (Li et al., 2009). For each candidate variant, information for all reads aligning to that position (e.g. base call, quality, coverage), across all samples, were aggregated into a data structure for local filtering and analysis. Colonies were omitted from further analysis if less than 90% of their reads were assigned to *Cutibacterium acnes* according to bracken (v2.5) (Lu et al., 2017; Wood et al., 2019) with the standard Univec database including all RefSeq genomes (153 of an initial 1546 colonies), if they had a median coverage below 10 across candidate variant positions (283 of 1393 colonies remaining), or if they had a major allele frequency below 0.65 for over 1% of variant positions with coverage greater or equal to 4 reads (50 colonies of 1110 remaining). In all, these filters retained 1080 colonies.

We filtered candidate SNVs using publicly available code (see Data Availability) similar to that previously published (Lieberman et al., 2014). Basecalls were marked as ambiguous if the FQ score produced by samtools was above −30, the coverage per strand was below 3, the major allele frequency was below 0.9, or more than 50% of reads supported indels. Remaining variant positions were discarded for clustering analysis if no unmasked polymorphisms remained. In addition, all SNVs in regions of the reference genome with homology to *C. acnes* plasmids were removed (see section on *C. acnes* plasmids). These SNV calls were used to calculate pairwise distances between colonies, equal to the number of positions where both colonies had non-ambiguous base calls and where the base calls differed. This distance matrix was used as input to clustering algorithm DBSCAN, using a distance threshold of 35 SNVs and a minimum cluster size of 3 (Figure S1). Clusters with a mean pairwise distance of below 80 SNVs were allowed to merge together (this allowed the hypermutator colonies to be part of Lineage 1a; see Figure S4).

Some colonies showed evidence of non-purity at this step, with mixed alleles at positions that distinguished colonies within the same initial cluster from each other. This nonpurity could have emerged during initial sample collection (no attempt was made to purify colonies into isolates before sequencing), during sample processing (all samples were processed in 96 well plates), or due to index hopping. Thus, after performing initial clustering, we removed colonies with a mean major allele frequency below 0.95 across within-cluster SNVs (variant positions that had base calls in at least 67% of colonies and with a median coverage of at least 10) for which the colony had sufficient coverage (greater or equal to 8 reads). Clustering and SNV identification were then repeated iteratively until no colonies with evidence of impurity remained, first by restricting clustering to colonies from the same subject only, and then by allowing clustering across subjects and allowing cluster merging (106 colonies were removed during this step). Finally, there were 7 colonies that clustered with Subject 1 despite originating from other subjects; these were removed due to suspected contamination, since Subject 1 was involved in sample acquisition and processing. In all, 947 high quality colonies and 53 clusters remained and were used in subsequent analysis (Tables S1, S2, and S4).

### Classification of lineages into *C. acnes* strains types

We used lineage-specific assemblies (see Mobile element analysis) to identify the global strain-types, using the previously described SLST scheme (Scholz et al., 2014). We used BLAST to compare known SLST types to custom BLAST databases created from lineage assemblies. Some lineages had no exact matches, indicating a new SLST. In this case, we classified the lineage by the super-SLST level (e.g. “A” for SLST “A1”), based on SLST with the best alignment (blastn with default parameters; highest bit score for alignment lengths greater or equal to 480 bp) (Altschul et al., 1990; Camacho et al., 2009). The super-SLST for each lineage is available in Table S4.

### SNV calling and evolutionary inference

In order to determine SNV positions within each lineage, basecalling was repeated using the following process: first, basecalls were marked as ambiguous if the FQ score produced by samtools was above −30, the coverage per strand was below 3, the major allele frequency was below 0.75, or more than 25% of reads supported indels; second, genomic positions with a median coverage below 12 reads across samples or where at least 34% of basecalls were ambiguous across samples were omitted. In addition, to remove variants that emerged from recombination or other complex events, we identified SNVs that were less then 500 bases apart and for which the correlation of non-ancestral allele frequencies (see below) across colonies within a lineage exceeded 0.75 (Table S7); these positions, as well as regions on the reference genome with homology to plasmids (see *C. acnes* plasmid analysis), were removed from downstream analysis.

All remaining genomic positions that passed these strict filters and retained two non-ambiguous alleles were considered SNV positions and were investigated across samples. In order to have genotypes for as many colonies as possible at these SNV positions, including ones with low coverage, basecalls were repopulated from the raw data and only marked as ambiguous only if the coverage per strand was below 1, the total coverage below 3, or the major allele frequency below 0.67. Details on SNVs detected in each lineage are available in Table S5.

Phylogenetic reconstruction was done using dnapars from PHYLIP v3.69 (Fenselstein, 2005). Trees were rooted using the ancestral allele as determined below. Example trees are shown in Figure 4 and Figures S4-6. Ancestral alleles were determined by using the most closely related lineage from a different subject (as measured by mean pairwise distance between colonies belonging to different lineages) as an outgroup: the ancestral allele was taken as the most common allele across 10 random colonies from the outgroup (or fewer colonies if the outgroup lineage contained less than 10 colonies). If outgroup colonies did not have any calls at that position, then the reference genome was used as the ancestral allele.

Phylogenetic reconstruction across lineages (Figure 2C) was performed using the inferred ancestral genotype of each lineage (for positions that did not vary within the lineage, the ancestral genotype was taken as the basecall across non-ambiguous samples; for positions that did vary within the lineage, the ancestral genotype was determined from an outgroup as described above). A parsimony tree was generated using dnapars as above, using variable positions with basecalls in greater than 10% of lineage ancestors. The tree is midpoint-rooted.

### Calculation of distances to MRCAs

To understand the evolutionary history of bacteria within and between pores, we calculated values of dMRCA (distance to most recent common ancestor) for sets of colonies (Figures 4 and 5). For vertically evolving organisms, this value has more interpretability than other metrics of diversity (e.g. average pairwise difference), representing the relative time since the set of organisms under consideration had a single-celled ancestor. In addition, dMRCA is robust to unequal sampling depth between clades on a phylogeny.

For each calculation of dMRCA, we inferred the genotype of the MRCA by assuming that, for each variable genomic position within the set of colonies, the ancestral allele was equal to that defined for the lineage ancestor (see SNV calling and evolutionary inference). We define the dMRCA for each pore as the mean of the number of SNVs distinguishing each colony from the pore MRCA. We exclude multipore samples as well as pore samples with only a single colony from calculations of intrapore dMRCAs. We define the interpore dMRCA for a pair of pores as the mean number of SNVs distinguishing the MRCA of each of the two pores and interpore MRCA. The genetic distances between pores reported in Figure 5C refer to the number of SNVs differing between the inferred ancestors of a given pair of pores.

### Parallel evolution analysis

In order to search for genes with evidence of mutational enrichment, we first counted how many times each gene was mutated (*m_i_*). We then computed the probability of observing ≥ *m_i_* mutations according to a Poisson distribution with λ = *Mp_i_*, where *M* is the total number of mutations observed on coding regions and *p_i_* is the expected probability that a random mutation lands that gene (taking into account gene length, codon distribution, and observed mutational spectrum). This analysis masked all regions of the reference genome with homology to *C. acnes* plasmids (see *C. acnes* plasmid analysis). To account for multiple hypotheses, we performed the Benjamini-Hochberg procedure (treating each unmasked gene on the genome as a hypothesis). We find no compelling evidence of parallel evolution when considering all *de novo* mutations or mutations at the intrasubject or intralineage levels (Figure S10).

In order to look for signs of adaptation, we computed dN/dS, the ratio of nonsynonymous mutations to synonymous mutations relative to a neutral model. Observed mutations were called as nonsynonymous (N) or synonymous (S) according to the reading frames in the annotated reference genome; in the event that there was an ancestral mutation (fixed in all colonies in a lineage) that differed from the reference genome, the basecall at that position was considered when determining if a SNV on that codon was N or S. Our neutral model was used to assess the expected N/S ratio, based on the observed mutational spectrum and the codon distribution of each gene. Figure 5B shows a summary of this analysis, and Figure S10 shows an extended version that considers mutations by subject, lineage, mutational age, and gene function. All results were consistent with neutrality or with weak purifying selection. We note that one limitation of this study is that it focused on ongoing evolution and would not capture any potential adaptive sweeps that occurred in the past, for example, immediately after a strain colonizes an individual.

### Mobile element analysis

To systematically identify gains and losses within each lineage, we constructed pan-genomes for each lineage. Only colonies with ≥ 95% of reads assigned to *C. acnes* at the species level by bracken were included (see Clustering colonies into lineages). For each lineage, we then concatenated up to 250,000 reads from each member colony and assembled a reference genome with SPAdes (v3.13, careful mode) (Bankevich et al., 2012) and a minimum contig length of 500 bp. We then aligned reads from each member colony to its assembled pangenome (see SNV calling and evolutionary inference).

We then looked for genomic regions that were missing from some, but not all, colonies in a lineage. We identified candidate mobile elements as continuous regions over 500 bp with a copy number (relative to the rest of the genome) below 0.25x in a given colony. We also considered each contig from the assembled genome as a candidate mobile element region. We then filtered these candidate regions, requiring a mean copy number (relative to the rest of the genome) less than or equal to 0.15x and mean coverage of less than or equal to 2.5 reads; we also required that the region to have strong support in at least one other colony (mean copy number greater or equal to 0.85x). Regions with homology to *C. acnes* plasmids were masked (see next section). We merged all overlapping regions found in colonies from the same lineage, and these regions are reported in Table S3.

### *C. acnes* plasmid analysis

During the mobile element analysis, we noted the presence of gain/loss regions with homology to known *C. acnes* plasmids (NCBI CP003294 and CP017041) (Brüggemann et al., 2012; Kasimatis et al., 2013). We also suspected that additional gain/loss regions belonged to plasmids because they had similar coverage patterns across samples to known plasmid regions. To better understand plasmid architecture and identify additional plasmid gene content, we performed long read sequencing for five isolates, which cover diverse genotypes on Subject 1: subj-1_scrape-439_col-5, subj-1_scrape-440_col-3, subj-1_scrape-441_col-3, subj-1_scrape-442_col-6, and subj-1_scrape-443_col-6. We used the Qiagen High-Molecular Weight Genomic DNA Kit (Catalog #67563) following the protocol recommended for gram-positive bacteria, but with increased lysozyme as above (see Culturing and single-colony sequencing). MIT’s BioMicroCenter performed size selection with SPRI beads to remove fragments below 10 kbp, library preparation with Oxford Nanopore kits EXP-NBD104 and SQK-LSK109, and sequencing on a R9 PromethION flow cell over 72 hours. Long reads were filtered using filtlong (v0.2.0, -- min_length 20000 --keep_percent 99 --target_bases 500000000) (Wick, 2018), and hybrid assemblies were generated using unicycler (v0.4.8, default parameters) (Wick et al., 2017). Scaffolds with homology (blastn, default parameters, total alignment lengths over 2,000 bp) to known *C. acnes* plasmids were designated as plasmid scaffolds and ranged in length from 16,360bp to 72,583bp (two contigs from isolate subj-1_scrape-440_col-3 were concatenated; all others had a single plasmid contig).

In order to determine which colonies in our dataset had evidence of plasmid presence, we aligned short reads to these five plasmid scaffolds using the same procedure for alignments to the *C. acnes* reference genome. A colony was deemed as having a plasmid if it had a copy number over 0.33x (relative to rest of genome) across at least 90% of at least one of the plasmid scaffolds. Plasmid presence/absence is available in Table S2 and indicated on lineage trees in Figures S4-6. To see how the plasmids in our dataset were related to each other, we generated a phylogenetic tree comparing a region common to as many plasmids as possible. We used alignments to the plasmid scaffold generated from isolate subj-1_scrape-441_col-3 (this scaffold had the most lineages with at least one positive colony). We then masked positions where fewer than 67% of plasmid-positive colonies had a copy number over 0.75 and removed colonies that had a copy number of less than 0.75 over fewer than 75% of these positions (this maintained 215 of 291 plasmid-positive colonies). Basecalls were marked as ambiguous if the quality was below 30, the coverage per strand was below 3, or the major allele frequency was below 0.67. This retained 769 variable positions, which were used to generate a parsimony tree using the same procedure as for lineage trees (Figure S7).

In order to avoid calling SNVs on mobile elements for genome focused analyses, we masked plasmid regions on the reference genome where there was an alignment to one of our plasmid scaffolds or to known plasmid genotypes CP003294 and CP017041 (blastn using default parameters with a minimum alignment length of 200 bp and a maximum e-value of 0.001). In our analysis of gain/loss regions, we additionally masked any contig for which these alignments covered over half of the contig positions.

### *Cutibacterium granulosum* analysis

There were 50 colonies for which greater or equal to 75% of reads were assigned as *Cutibacterium granulosum* according to bracken (see Clustering colonies into lineages). In order to characterize the within-species *C. granulosum* diversity, we used an alignment-based approach following the same procedure as above, but with *C. granulosum* NCTC 11865 (RefSeq NZ_LT906441.1) as the reference genome. Colonies were removed from the analysis if less than 72% of reads aligned to the reference genome (7 colonies) or if they had a mean coverage of 5x or below across candidate variant positions (1 colony). Basecalls were marked as ambiguous if the FQ score produced by samtools was above −30, the coverage per strand was below 3, or the major allele frequency was below .75. Remaining variant positions were discarded if 34% or more of all colonies were called as ambiguous, if the median coverage across all colonies at that position was below 12, or if no polymorphisms remained. Any colonies for which greater than 30% of variant positions were marked as ambiguous at this stage were removed (this removed 3 colonies). In all, these filters retained ~90,000 variable positions across 39 colonies. Basecalls were repopulated from the raw data and only marked as ambiguous only if the FQ score was above −30, the coverage was below 5, or the major allele frequency was below .67. A parsimony tree (Figure S14, left panel) was generated using the same process as for *C. acnes*.

We identified three pores (Subject 1, pore 17; Subject 1, pore 18; Subject 2, pore 87) for which there were multiple *C. granulosum* colonies and for which these colonies were monophyletic on the tree constructed above. For each case, we assembled a genome using the same procedure as for *C. acnes* lineages, using reads from all colonies from that pore. We then aligned reads from each colony (including all colonies from that pore and any additional colonies within 100 SNVs according to the above tree) onto its pore-specific assembled genome and called SNVs using the same filters as above in order to generate parsimony trees (Figure S14, right panels).

### 16S amplicon sequencing

Samples collected for community profiling were collected in QuickExtract buffer (see Human subjects and sample collection). After streaking for single colonies, the remainder of samples were lysed by adding 1 ul of ReadyLyse (EpiCentre) and incubating at room temperature for 12 hours. A 1 uL aliquot was used to amplify the V1-V3 region using HiFi HotStart ReadyMix (KAPA BioSystems) and the Illumina PCR protocol. A spike of genomic DNA from *C*aul*obacter crecentus*, a species typically found in freshwater, was included in each PCR reaction to estimate the number of unique sequencing reads. Samples were cleaned and pooled as in (Baym et al., 2015). Samples were sequenced on an Illumina MiSeq (300 PE) to an average read depth of ~16,500.

In order to classify amplicon sequence variants (ASV) on a species level, a classifier was built using the V1-V3 region of the raw sequences and taxonomy of the SILVA database (version 132) (Quast et al., 2013), with taxonomically-mislabeled sequences identified by the phylogeny-aware pipeline SATIVA (Kozlov et al., 2016) either corrected or removed. *Staphylococcus* species were specifically filtered by the methods presented in (Khadka et al., 2021). The genuses *Cutibacterium*, *Acidipropionibacterium*, *Pseudopropionibacterium*, *Propionibacterium*, and the *Corynebacteriaceae* and chronically-mislabelled *Neisseriaceae* families were also cleaned by the following filters: (i) sequences with incorrect higher taxonomic classes (ex. a species with the family Corynebacteriaceae but the genus Cutibacterium) were removed, (ii) sequences missing a species classification or assigned to non-species taxa (ex. Corynebacterium sp.) were removed, (iii) species with >60% similarity with other taxa were relabeled as a specific “taxa cluster”, (iv) taxonomically mislabeled sequences identified using SATIVA with greater than 90% confidence were relabeled and sequences with below 90% confidence removed. To reduce computational load, each family or genuses within the same family were grouped together in independent SATIVA runs. This removed about 2% of sequences from each group. This database was then used to train a naive Bayes classifier in QIIME2 (2020.01).

QIIME2 was used to process and classify 16S reads, using cutadapt and DADA2 (Bolyen et al., 2018; Callahan et al., 2016; Martin, 2011). In order to visualize species diversity, all spike-in sequences, unclassified reads, and reads with only a domain-level classification were removed from further analysis (Figure S9).

## DATA AND SOFTWARE AVAILABILITY

All raw sequencing data will be deposited to the SRA prior to publication under submission number SUB9587388. Data analysis code and genome assemblies will be available at https://github.com/arolynconwill/cacnes_biogeo.

## AUTHOR CONTRIBUTIONS

T.D.L. designed the study within input from E.J.A., T.D.L., and A.K. collected samples from human subjects. T.D.L. and J.S.B. prepared samples for whole genome sequencing. A.C. and T.D.L. analyzed the data. A.D.T. performed growth curve experiments. A.K., A.P., and T.D.L. collected and analyzed 16S amplicon sequencing data. R.D. and A.C. prepared samples for long read sequencing and analyzed mobile genetic elements. E.J.A. and T.D.L. secured funding and materials. A.C. and T.D.L. wrote the manuscript with input from all authors.

## ACKNOWLEDGMENTS

We thank Sarah Bi, Sean Kearny, Sean Gibbons, Allison Perrotta, Claire Duvallet, Tucker Lynn, and all Lieberman Lab members for experimental assistance and helpful discussions, the MIT BioMicroCenter for assistance in sequencing, and Oskar Hallatschek for helpful discussions. We thank Vicki Mountain and Scott Olesen for assistance with IRB protocols. This work was funded by grants from the Broad Institute (to E.J.A.), the Smith Family Foundation (to T.D.L.), the MIT Center for Microbiome Informatics and Therapeutics (to T.D.L.), and the National Institutes of Health (1DP2GM140922-01 to T.D.L.).

## References

Adamson, A.S., and Lipoff, J.B. (2021). Reconsidering Named Honorifics in Medicine—the Troubling Legacy of Dermatologist Albert Kligman. Jama Dermatol 157, 153–155.

Altschul, S.F., Gish, W., Miller, W., Myers, E.W., and Lipman, D.J. (1990). Basic local alignment search tool. J Mol Biol 215, 403–410.

Bankevich, A., Nurk, S., Antipov, D., Gurevich, A.A., Dvorkin, M., Kulikov, A.S., Lesin, V.M., Nikolenko, S.I., Pham, S., Prjibelski, A.D., et al. (2012). SPAdes: A New Genome Assembly Algorithm and Its Applications to Single-Cell Sequencing. J Comput Biol 19, 455–477.

Barrick, J.E., and Lenski, R.E. (2013). Genome dynamics during experimental evolution. Nat Rev Genet 14, 827–839.

Baym, M., Kryazhimskiy, S., Lieberman, T.D., Chung, H., Desai, M.M., and Kishony, R. (2015). Inexpensive Multiplexed Library Preparation for Megabase-Sized Genomes. Plos One 10, e0128036.

Bolyen, E., Rideout, J.R., Dillon, M.R., Bokulich, N.A., Abnet, C., Al-Ghalith, G.A., Alexander, H., Alm, E.J., Arumugam, M., Asnicar, F., et al. (2018). QIIME 2: Reproducible, interactive, scalable, and extensible microbiome data science. Peerj Prepr 6, e27295v2.

Brüggemann, H., Henne, A., Hoster, F., Liesegang, H., Wiezer, A., Strittmatter, A., Hujer, S., Dürre, P., and Gottschalk, G. (2004). The Complete Genome Sequence of Propionibacterium Acnes, a Commensal of Human Skin. Science 305, 671.

Brüggemann, H., Lomholt, H.B., Tettelin, H., and Kilian, M. (2012). CRISPR/cas Loci of Type II Propionibacterium acnes Confer Immunity against Acquisition of Mobile Elements Present in Type I P. acnes. Plos One 7, e34171.

Brzuszkiewicz, E., Weiner, J., Wollherr, A., Thürmer, A., Hüpeden, J., Lomholt, H.B., Kilian, M., Gottschalk, G., Daniel, R., Mollenkopf, H.-J., et al. (2011). Comparative Genomics and Transcriptomics of Propionibacterium acnes. Plos One 6, e21581.

Butcher, E., and Coonin, A. (1949). The physical properties of human sebum. J Investigative Dermatology 12, 249–254.

Byrd, A.L., Belkaid, Y., and Segre, J.A. (2018). The human skin microbiome. Nat Rev Microbiol 16, 143–155.

Callahan, B.J., McMurdie, P.J., Rosen, M.J., Han, A.W., Johnson, A.J.A., and Holmes, S.P. (2016). DADA2: High-resolution sample inference from Illumina amplicon data. Nat Methods 13, 581–583.

Camacho, C., Coulouris, G., Avagyan, V., Ma, N., Papadopoulos, J., Bealer, K., and Madden, T.L. (2009). BLAST+: architecture and applications. Bmc Bioinformatics 10, 421.

Castañeda-García, A., Martín-Blecua, I., Cebrián-Sastre, E., Chiner-Oms, A., Torres-Puente, M., Comas, I., and Blázquez, J. (2020). Specificity and mutagenesis bias of the mycobacterial alternative mismatch repair analyzed by mutation accumulation studies. Sci Adv 6, eaay4453.

Chung, H., Lieberman, T.D., Vargas, S.O., Flett, K.B., McAdam, A.J., Priebe, G.P., and Kishony, R. (2017). Global and local selection acting on the pathogen Stenotrophomonas maltophilia in the human lung. Nat Commun 8, 14078.

Claesen, J., Spagnolo, J.B., Ramos, S.F., Kurita, K.L., Byrd, A.L., Aksenov, A.A., Melnik, A.V., Wong, W.R., Wang, S., Hernandez, R.D., et al. (2020). A Cutibacterium acnes antibiotic modulates human skin microbiota composition in hair follicles. Sci Transl Med 12, eaay5445.

Costello, E.K., Lauber, C.L., Hamady, M., Fierer, N., Gordon, J.I., and Knight, R. (2009). Bacterial Community Variation in Human Body Habitats Across Space and Time. Science 326, 1694–1697.

Cove, J.H., Holland, K.T., and Cunliffe, W.J. (1983). Effects of Oxygen Concentration on Biomass Production, Maximum Specific Growth Rate and Extracellular Enzyme Production by Three Species of Cutaneous Propionibacteria Grown in Continuous Culture. Microbiology 129, 3327–3334.

Coyte, K.Z., Schluter, J., and Foster, K.R. (2015). The ecology of the microbiome: Networks, competition, and stability. Science 350, 663–666.

Didelot, X., Walker, A.S., Peto, T.E., Crook, D.W., and Wilson, D.J. (2016). Within-host evolution of bacterial pathogens. Nat Rev Microbiol 14, 150–162.

Dréno, B., Pécastaings, S., Corvec, S., Veraldi, S., Khammari, A., and Roques, C. (2018). Cutibacterium acnes (Propionibacterium acnes) and acne vulgaris: a brief look at the latest updates. J Eur Acad Dermatol 32, 5–14.

Fenselstein, J. (2005). PHYLIP (Phylogeny Inference Package) version 3.6 (Distributed by the author: Department of Genome Sciences, University of Washington, Seattle).

Ferreiro, A., Crook, N., Gasparrini, A.J., and Dantas, G. (2018). Multiscale Evolutionary Dynamics of Host-Associated Microbiomes. Cell 172, 1216–1227.

Fitz-Gibbon, S., Tomida, S., Chiu, B.-H., Nguyen, L., Du, C., Liu, M., Elashoff, D., Erfe, M.C., Loncaric, A., Kim, J., et al. (2013). Propionibacterium acnes Strain Populations in the Human Skin Microbiome Associated with Acne. J Invest Dermatol 133, 2152–2160.

Flowers, L., and Grice, E.A. (2020). The Skin Microbiota: Balancing Risk and Reward. Cell Host Microbe 28, 190–200.

Foster, K.R., Schluter, J., Coyte, K.Z., and Rakoff-Nahoum, S. (2017). The evolution of the host microbiome as an ecosystem on a leash. Nature 548, 43–51.

Garud, N.R., Good, B.H., Hallatschek, O., and Pollard, K.S. (2019). Evolutionary dynamics of bacteria in the gut microbiome within and across hosts. Plos Biol 17, e3000102.

Ghalayini, M., Launay, A., Bridier-Nahmias, A., Clermont, O., Denamur, E., Lescat, M., and Tenaillon, O. (2018). Evolution of a Dominant Natural Isolate of Escherichia coli in the Human Gut over the Course of a Year Suggests a Neutral Evolution with Reduced Effective Population Size. Appl Environ Microb 84, e02377–17.

Grice, E.A., and Segre, J.A. (2011). The skin microbiome. Nat Rev Microbiol 9, 244–253.

Hall, J.B., Cong, Z., Imamura-Kawasawa, Y., Kidd, B.A., Dudley, J.T., Thiboutot, D.M., and Nelson, A.M. (2018). Isolation and Identification of the Follicular Microbiome: Implications for Acne Research. J Invest Dermatol 138, 2033–2040.

Hartl, D.L., and Clark, A.G. (2006). Principles of Population Genetics (Oxford University Press).

Hecht, A.L., Casterline, B.W., Earley, Z.M., Goo, Y.A., Goodlett, D.R., and Wardenburg, J.B. (2016). Strain competition restricts colonization of an enteric pathogen and prevents colitis. Embo Rep 17, 1281–1291.

Ishino, S., Skouloubris, S., Kudo, H., l’Hermitte-Stead, C., Es-Sadik, A., Lambry, J.-C., Ishino, Y., and Myllykallio, H. (2018). Activation of the mismatch-specific endonuclease EndoMS/NucS by the replication clamp is required for high fidelity DNA replication. Nucleic Acids Res 46, gky460-.

Jahns, A.C., and Alexeyev, O.A. (2014). Three dimensional distribution of Propionibacterium acnes biofilms in human skin. Exp Dermatol 23, 687–689.

Jorth, P., Staudinger, B.J., Wu, X., Hisert, K.B., Hayden, H., Garudathri, J., Harding, C.L., Radey, M.C., Rezayat, A., Bautista, G., et al. (2015). Regional Isolation Drives Bacterial Diversification within Cystic Fibrosis Lungs. Cell Host Microbe 18, 307–319.

Joshi, N., and Fass, J. (2011). Sickle: A sliding-window, adaptive, quality-based trimming tool for FastQ files [software].

Kasimatis, G., Fitz-Gibbon, S., Tomida, S., Wong, M., and Li, H. (2013). Analysis of Complete Genomes of Propionibacterium acnes Reveals a Novel Plasmid and Increased Pseudogenes in an Acne Associated Strain. Biomed Res Int 2013, 1–11.

Kerr, B., Riley, M.A., Feldman, M.W., and Bohannan, B.J.M. (2002). Local dispersal promotes biodiversity in a real-life game of rock–paper–scissors. Nature 418, 171–174.

Khadka, V.D., Key, F.M., Romo-González, C., Martínez-Gayosso, A., Campos-Cabrera, B.L., Gerónimo-Gallegos, A., Lynn, T.C., Durán-McKinster, C., Coria-Jiménez, R., Lieberman, T.D., et al. (2021). A randomised longitudinal study of the skin microbiome during treatment for atopic dermatitis in children. In Review.

Koskella, B., Hall, L.J., and Metcalf, C.J.E. (2017). The microbiome beyond the horizon of ecological and evolutionary theory. Nat Ecol Evol 1, 1606–1615.

Kozlov, A.M., Zhang, J., Yilmaz, P., Glöckner, F.O., and Stamatakis, A. (2016). Phylogeny-aware identification and correction of taxonomically mislabeled sequences. Nucleic Acids Res 44, 5022–5033.

Ladau, J., and Eloe-Fadrosh, E.A. (2019). Spatial, Temporal, and Phylogenetic Scales of Microbial Ecology. Trends Microbiol 27, 662–669.

Langmead, B., Trapnell, C., Pop, M., and Salzberg, S.L. (2009). Ultrafast and memory-efficient alignment of short DNA sequences to the human genome. Genome Biol 10, R25.

LeClerc, J.E., Li, B., Payne, W.L., and Cebula, T.A. (1996). High Mutation Frequencies Among Escherichia coli and Salmonella Pathogens. Science 274, 1208–1211.

Leeming, J.P., Holland, K.T., and Cunliffe, W.J. (1984). The Microbial Ecology of Pilosebaceous Units Isolated from Human Skin. Microbiology+ 130, 803–807.

Li, H., Handsaker, B., Wysoker, A., Fennell, T., Ruan, J., Homer, N., Marth, G., Abecasis, G., Durbin, R., and Subgroup, 1000 Genome Project Data Processing (2009). The Sequence Alignment/Map format and SAMtools. Bioinformatics 25, 2078–2079.

Lieberman, E., Hauert, C., and Nowak, M.A. (2005). Evolutionary dynamics on graphs. Nature 433, 312–316.

Lieberman, T.D., Michel, J.-B., Aingaran, M., Potter-Bynoe, G., Roux, D., Davis, M.R., Skurnik, D., Leiby, N., LiPuma, J.J., Goldberg, J.B., et al. (2011). Parallel bacterial evolution within multiple patients identifies candidate pathogenicity genes. Nat Genet 43, 1275–1280.

Lieberman, T.D., Flett, K.B., Yelin, I., Martin, T.R., McAdam, A.J., Priebe, G.P., and Kishony, R. (2014). Genetic variation of a bacterial pathogen within individuals with cystic fibrosis provides a record of selective pressures. Nat Genet 46, 82–87.

Lieberman, T.D., Wilson, D., Misra, R., Xiong, L.L., Moodley, P., Cohen, T., and Kishony, R. (2016). Genomic diversity in autopsy samples reveals within-host dissemination of HIV-associated Mycobacterium tuberculosis. Nat Med 22, 1470–1474.

Lomholt, H.B., Scholz, C.F.P., Brüggemann, H., Tettelin, H., and Kilian, M. (2017). A comparative study of Cutibacterium (Propionibacterium) acnes clones from acne patients and healthy controls. Anaerobe 47, 57–63.

Lourenço, M., Chaffringeon, L., Lamy-Besnier, Q., Pédron, T., Campagne, P., Eberl, C., Bérard, M., Stecher, B., Debarbieux, L., and Sordi, L.D. (2020). The Spatial Heterogeneity of the Gut Limits Predation and Fosters Coexistence of Bacteria and Bacteriophages. Cell Host Microbe 28, 390–401.e5.

Lu, J., Breitwieser, F.P., Thielen, P., and Salzberg, S.L. (2017). Bracken: estimating species abundance in metagenomics data. Peerj Comput Sci 3, e104.

Mak, T.N., Schmid, M., Brzuszkiewicz, E., Zeng, G., Meyer, R., Sfanos, K.S., Brinkmann, V., Meyer, T.F., and Brüggemann, H. (2013). Comparative genomics reveals distinct host-interacting traits of three major human-associated propionibacteria. Bmc Genomics 14, 640.

Martin, M. (2011). Cutadapt removes adapter sequences from high-throughput sequencing reads. Embnet J 17, 10–12.

McLaughlin, J., Watterson, S., Layton, A.M., Bjourson, A.J., Barnard, E., and McDowell, A. (2019). Propionibacterium acnes and Acne Vulgaris: New Insights from the Integration of Population Genetic, Multi-Omic, Biochemical and Host-Microbe Studies. Microorg 7, 128.

Minegishi, K., Aikawa, C., Furukawa, A., Watanabe, T., Nakano, T., Ogura, Y., Ohtsubo, Y., Kurokawa, K., Hayashi, T., Maruyama, F., et al. (2013). Complete Genome Sequence of a Propionibacterium acnes Isolate from a Sarcoidosis Patient. Genome Announc 1, e00016–12.

Miskin, J.E., Farrell, A.M., Cunliffe, W.J., and Holland, K.T. (1997). Propionibacterium acnes, a resident of lipid-rich human skin, produces a 33 kDa extracellular lipase encoded by gehA. Microbiology+ 143, 1745–1755.

Mölder, F., Jablonski, K.P., Letcher, B., Hall, M.B., Tomkins-Tinch, C.H., Sochat, V., Forster, J., Lee, S., Twardziok, S.O., Kanitz, A., et al. (2021). Sustainable data analysis with Snakemake. F1000research 10, 33.

Oh, J., Byrd, A.L., Deming, C., Conlan, S., Barnabas, B., Blakesley, R., Bouffard, G., Brooks, S., Coleman, H., Dekhtyar, M., et al. (2014). Biogeography and individuality shape function in the human skin metagenome. Nature 514, 59–64.

Oh, J., Byrd, A.L., Park, M., Program, N.C.S., Kong, H.H., and Segre, J.A. (2016). Temporal Stability of the Human Skin Microbiome. Cell 165, 854–866.

Oliver, A. (2010). Mutators in cystic fibrosis chronic lung infection: Prevalence, mechanisms, and consequences for antimicrobial therapy. International Journal of Medical Microbiology 300, 563–572.

O’Neill, A.M., and Gallo, R.L. (2018). Host-microbiome interactions and recent progress into understanding the biology of acne vulgaris. Microbiome 6, 177.

Paetzold, B., Willis, J.R., Lima de, J.P., Knödlseder, N., Brüggemann, H., Quist, S.R., Gabaldón, T., and Güell, M. (2019). Skin microbiome modulation induced by probiotic solutions. Microbiome 7, 95.

Plewig, G. (1974). Follicular Keratinization. J Invest Dermatol 62, 308–315.

Plewig, G., Melnik, B., and Chen, W. (2019). Plewig and Kligman’;s Acne and Rosacea.

Poyet, M., Groussin, M., Gibbons, S.M., Avila-Pacheco, J., Jiang, X., Kearney, S.M., Perrotta, A.R., Berdy, B., Zhao, S., Lieberman, T.D., et al. (2019). A library of human gut bacterial isolates paired with longitudinal multiomics data enables mechanistic microbiome research. Nat Med 25, 1442–1452.

Quast, C., Pruesse, E., Yilmaz, P., Gerken, J., Schweer, T., Yarza, P., Peplies, J., and Glöckner, F.O. (2013). The SILVA ribosomal RNA gene database project: improved data processing and web-based tools. Nucleic Acids Res 41, D590–D596.

Rossum, T.V., Ferretti, P., Maistrenko, O.M., and Bork, P. (2020). Diversity within species: interpreting strains in microbiomes. Nat Rev Microbiol 18, 491–506.

Schmidt, C. (2020). Out of your skin. Nat Biotechnol 38, 392–397.

Scholz, C.F.P., Jensen, A., Lomholt, H.B., Brüggemann, H., and Kilian, M. (2014). A Novel High-Resolution Single Locus Sequence Typing Scheme for Mixed Populations of Propionibacterium acnes In Vivo. Plos One 9, e104199.

Scholz, C.F.P., Brüggemann, H., Lomholt, H.B., Tettelin, H., and Kilian, M. (2016). Genome stability of Propionibacterium acnes: a comprehensive study of indels and homopolymeric tracts. Sci Rep-Uk 6, 20662.

Schreck, C.F., Fusco, D., Karita, Y., Martis, S., Kayser, J., Duvernoy, M.-C., and Hallatschek, O. (2019). Impact of crowding on the diversity of expanding populations. Biorxiv 743534.

Sniegowski, P.D., Gerrish, P.J., and Lenski, R.E. (1997). Evolution of high mutation rates in experimental populations of E. coli. Nature 387, 703–705.

Tenaillon, O., Barrick, J.E., Ribeck, N., Deatherage, D.E., Blanchard, J.L., Dasgupta, A., Wu, G.C., Wielgoss, S., Cruveiller, S., Médigue, C., et al. (2016). Tempo and mode of genome evolution in a 50,000-generation experiment. Nature 536, 165–170.

Tomida, S., Nguyen, L., Chiu, B.-H., Liu, J., Sodergren, E., Weinstock, G.M., and Li, H. (2013). Pan-Genome and Comparative Genome Analyses of Propionibacterium acnes Reveal Its Genomic Diversity in the Healthy and Diseased Human Skin Microbiome. Mbio 4, e00003–13.

Tropini, C., Earle, K.A., Huang, K.C., and Sonnenburg, J.L. (2017). The Gut Microbiome: Connecting Spatial Organization to Function. Cell Host Microbe 21, 433–442.

Welch, J.L.M., Rossetti, B.J., Rieken, C.W., Dewhirst, F.E., and Borisy, G.G. (2016). Biogeography of a human oral microbiome at the micron scale. Proc National Acad Sci 113, E791–E800.

Whitaker, W.R., Shepherd, E.S., and Sonnenburg, J.L. (2017). Tunable Expression Tools Enable Single-Cell Strain Distinction in the Gut Microbiome. Cell 169, 538–546.e12.

Wick, R. (2018). Filtlong v0.2.0 (https://github.com/rrwick/Filtlong).

Wick, R.R., Judd, L.M., Gorrie, C.L., and Holt, K.E. (2017). Unicycler: Resolving bacterial genome assemblies from short and long sequencing reads. Plos Comput Biol 13, e1005595.

Wielgoss, S., Barrick, J.E., Tenaillon, O., Wiser, M.J., Dittmar, W.J., Cruveiller, S., Chane-Woon-Ming, B., Médigue, C., Lenski, R.E., and Schneider, D. (2013). Mutation rate dynamics in a bacterial population reflect tension between adaptation and genetic load. Proc National Acad Sci 110, 222–227.

Wiser, M.J., Ribeck, N., and Lenski, R.E. (2013). Long-Term Dynamics of Adaptation in Asexual Populations. Science 342, 1364–1367.

Wood, D.E., Lu, J., and Langmead, B. (2019). Improved metagenomic analysis with Kraken 2. Genome Biol 20, 257.

Zhao, S., Lieberman, T.D., Poyet, M., Kauffman, K.M., Gibbons, S.M., Groussin, M., Xavier, R.J., and Alm, E.J. (2019). Adaptive Evolution within Gut Microbiomes of Healthy People. Cell Host Microbe 25, 656–667.e8.

Zhou, W., Spoto, M., Hardy, R., Guan, C., Fleming, E., Larson, P.J., Brown, J.S., and Oh, J. (2020). Host-Specific Evolutionary and Transmission Dynamics Shape the Functional Diversification of Staphylococcus epidermidis in Human Skin. Cell 180, 454–470.e18.

