## Supplemental Figures for "Anatomy promotes neutral coexistence of strains in the human skin microbiome"

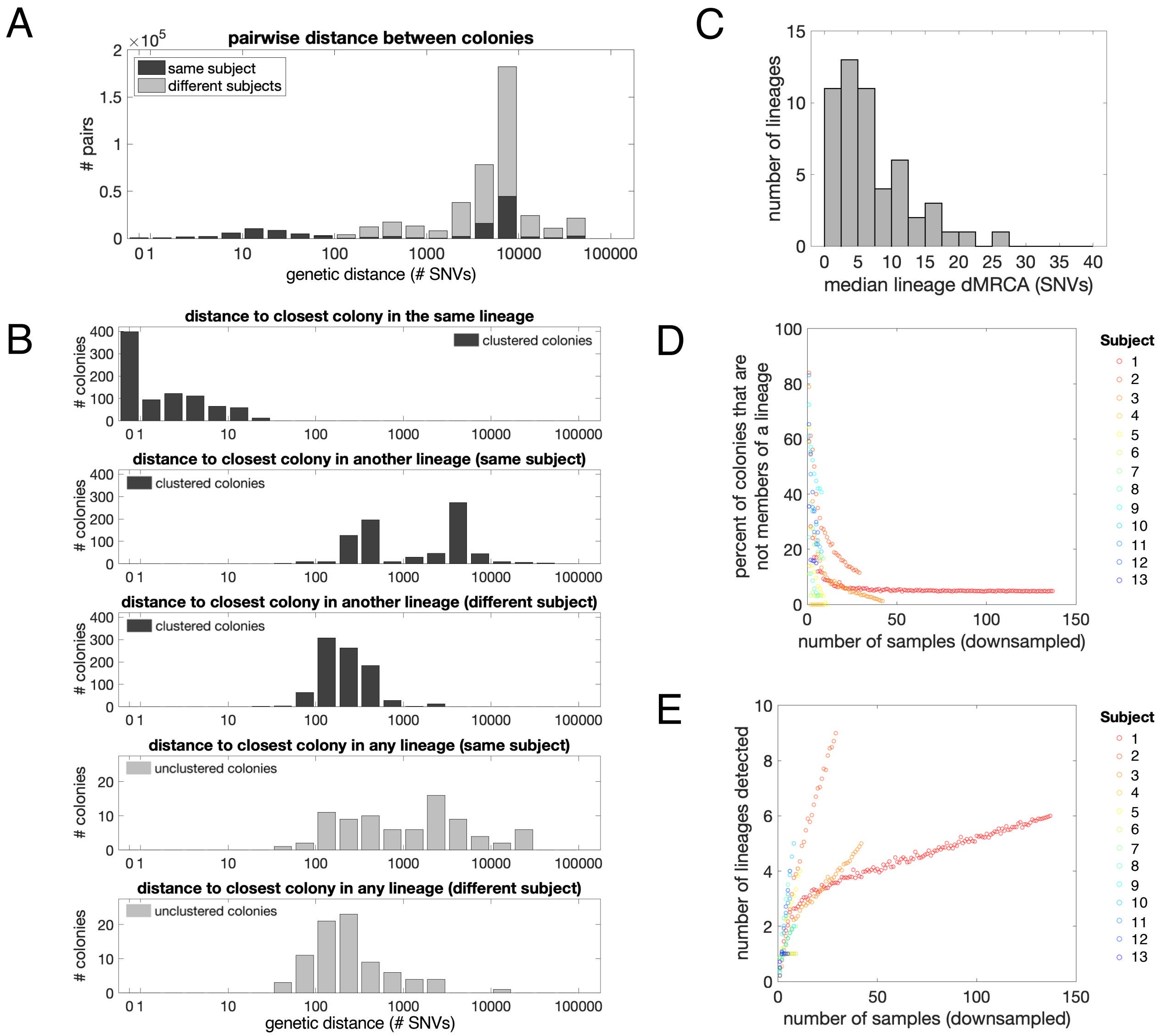


**Figure S1. Clustering distance thresholds and collector's curves.** (A) Histogram of the genetic distance (in single nucleotide variants) between pairs of colonies from the same subject vs different subjects. The bimodal distribution for same-subject pairs reflects that *C. acnes* genotypes found on an individual cluster into closely related lineages. (B) Histograms of the distance (in single nucleotide variants) to the nearest colony from various sets (see subpanel titles). The top three histograms consider clustered colonies (dark gray), while the bottom two histograms consider unclustered colonies (light gray). The minimum distance from a clustered colony to a lineage on another subject is generally lower than the minimum distance to another lineage on the same subject because the set of colonies belonging to other subjects is larger and thus contains representatives from a broader selection of strain types. In (C), we plot a histogram of the median distance between a lineage's inferred ancestor and the colonies belonging to that lineage. For each subject, we show collector's curves for (D) the percent of colonies that do not cluster with any lineage and (E) the number of lineages detected. Each dot represents the average over 100 downsampled sets; we downsample samples rather than colonies, as colonies from the same sample are more likely to be from the same lineage. We note 3 colonies are required to define a lineage, which results in a collector's curve with a steeper tail (E) than in a traditional collector's curve. The lack of saturation reflects the presence of low-abundance lineages and demonstrates the difficulty in estimating the true number of resident lineages.

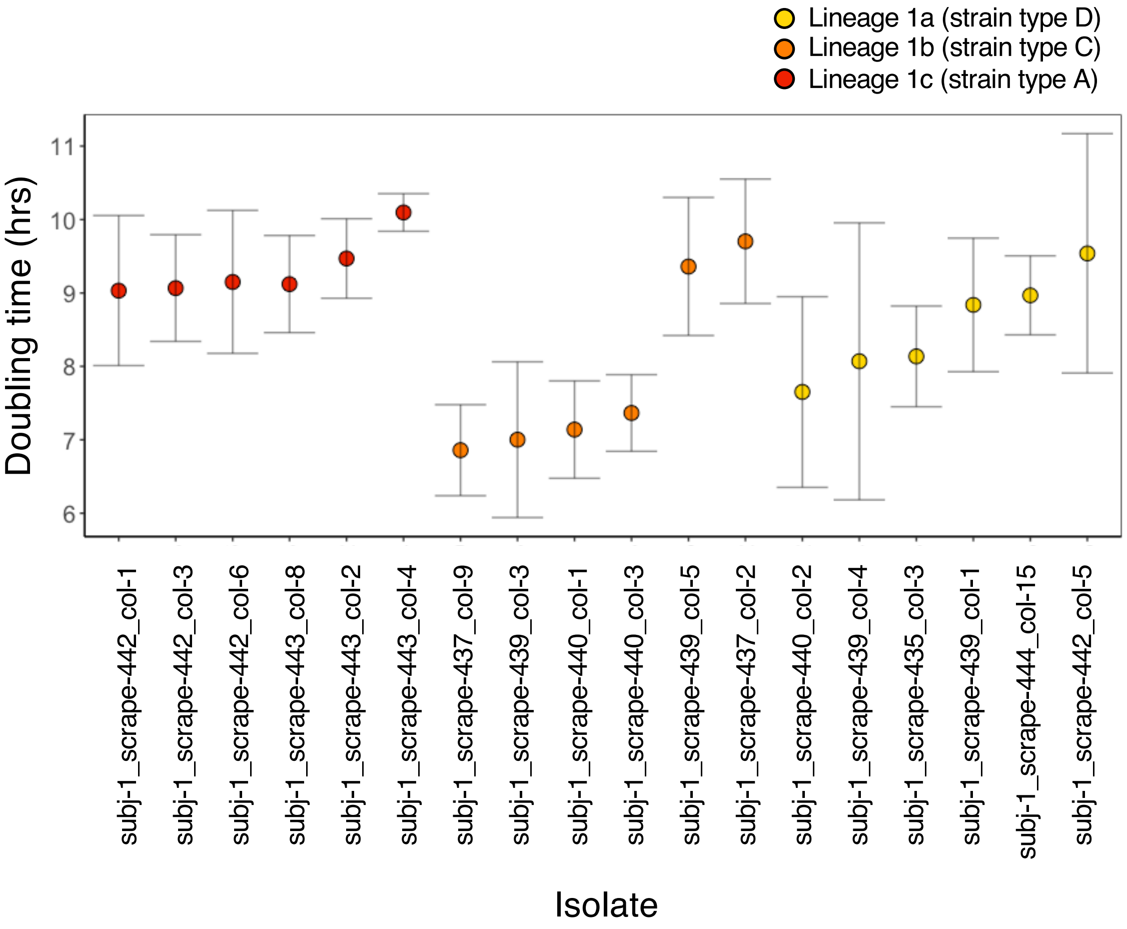


**Figure S2. Co-existing *C. acnes* genotypes have different doubling times *in vitro*.** All isolates tested were collected from Subject 1 at the same timepoint. Frozen stocks of isolates were revived on RCM plates (Oxoid CM0149) that were reduced overnight anaerobically at 33°C. For each strain, 3 independent replicates were started from 3 distinct colonies on the plate and grown to saturation over 4 days in deep-well plates in 300 uL of reduced RCM in anaerobic conditions at 33°C. From this, cultures for growth curves were inoculated by diluting 2 µl of saturated culture into 198 µl of RCM in microtiter plates. Growth curves were obtained inside a Tecan M Nano, taking readings of $OD_{595\ nm}$ every 15 minutes. Growth rates were obtained for each replicate by taking the slope from a linear regression of $\ln(OD)$ versus time during the period of exponential growth and then converted to doubling times. Each dot represents the mean observed doubling time among replicates for a given isolate, and error bars represent 95% CIs of the mean doubling time for each isolate (+/- 1.97 SEM). Color indicates phylogenetic grouping by lineage. The isolates do not all have the same doubling time (one way ANOVA, $p = 1.2\times10^{-13}$).

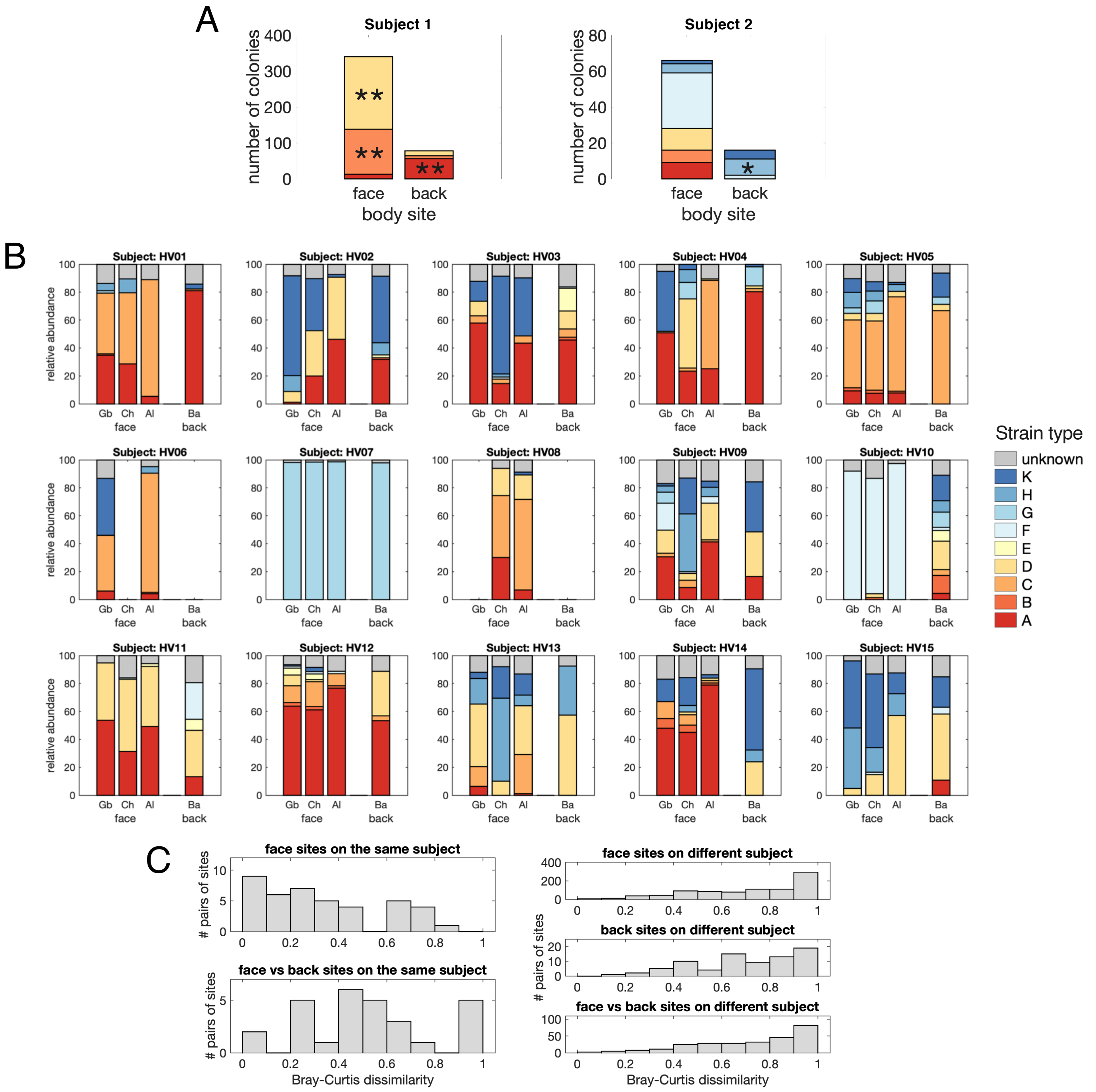


**Figure S3. Different strain enrichment patterns on the face vs the back on some subjects suggest a priority effect.** (A) Bar charts showing the number of colonies belonging to each strain type on the face vs back for subjects with at least 10 colonies on each site. Some strains are enriched on the face vs the back (**p*<0.01, ** *p*<0.001, binomial test with Bonferroni correction). (B,C) Reanalysis of published data by grouping relative abundances (from Supplemental Table 11 in Oh et al., 2014) at the strain level. Variation between face sites (Gb = glabella, Ch = cheek, Al = alar crease) and the back (Ba) differs between individual subjects. (C) Face sites on the same subject are more similar to each other than they are to the back (Bray-Curtis dissimilarity of strain type relative abundance; Wilcoxon rank sum test, *p*<0.01). However, back sites on different subjects and face sites on different subjects are not significantly more similar than face vs back sites on different subjects (Wilcoxon rank sum test). Since enrichment patterns are not conserved across subjects, these findings support a priority effect rather than generic back-adapted or face-adapted strains.

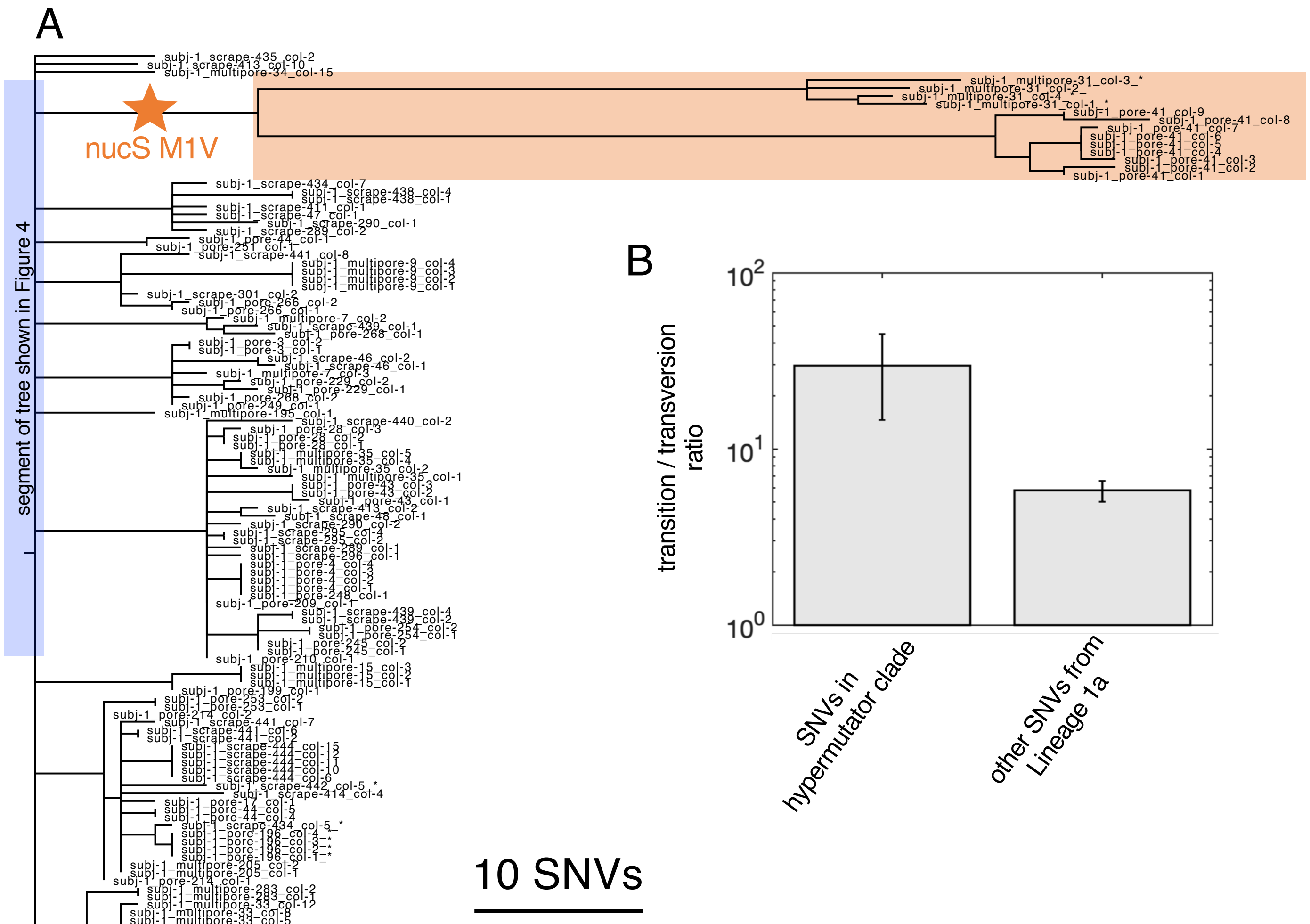


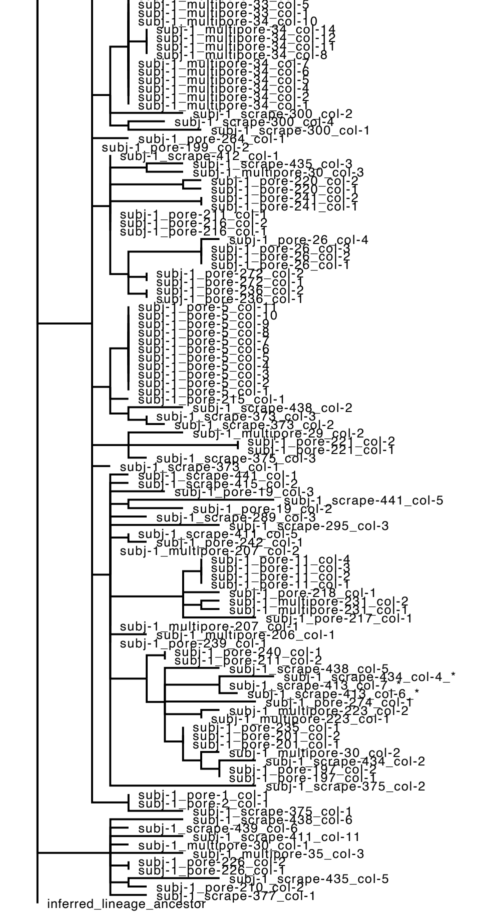


**Figure S4. Maximum parsimony phylogeny of Lineage 1a and evidence of hypermutation.** Full tree for Lineage 1a (blue bar to left side shows which portion of the tree was shown in Figure 4A). Tip labels indicate individual colonies, named by their subject and sample of origin; an asterisk in the colony name indicates the presence of a plasmid (see also Figure S7). Long branch lengths in the clade highlighted in orange suggest a hypermutator phenotype. In line with this, there is a nonsynonymous mutation in the start codon (M1V) of a gene implicated in DNA repair (endonuclease NucS, PAC1_RS05680) found in the branch leading to this clade. (B) SNVs within the hypermutator clade have a higher transition to transversion ratio than all mutations (p=.00015, one-sided Fisher exact test). Error bars represent 95% confidence intervals assuming binomial sampling. These findings are consistent with other work that has shown that a nonfunctional NucS gene results in a hypermutator phenotype with a enrichment in transition mutations (Castañeda-García et al., 2020; Ishino et al., 2018}. No other mutations on the NucS gene were detected in any lineage.

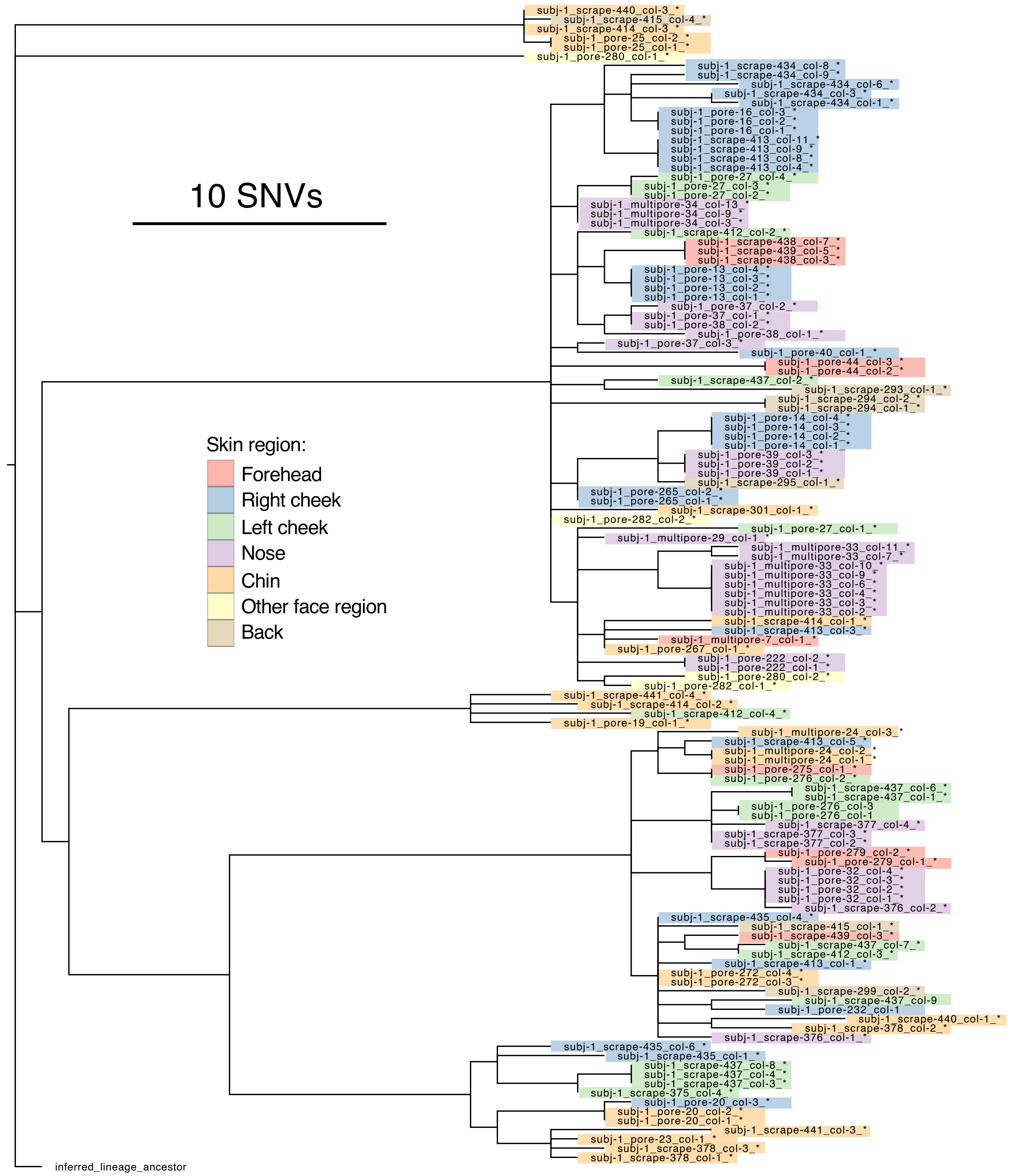


**Figure S5. Maximum parsimony phylogeny of Lineage 1b.** Full lineage tree of the second most abundant lineage on Subject 1, which also shows a pattern of low diversity within pores and pore-specific genotypes. Tip labels indicate individual colonies, named by their subject and sample of origin; an asterisk at the end of the colony name indicates the presence of a plasmid (see Figure S7). Long branches leading to subclades might reflect the transmission of multiple closely related genotypes to Subject 1 from a single source (e.g. a parent), or an expansion of particular genotypes following colonization (neutral or adaptive expansion). Highlight colors indicate skin region and show that closely related genotypes are not necessarily co-localized.

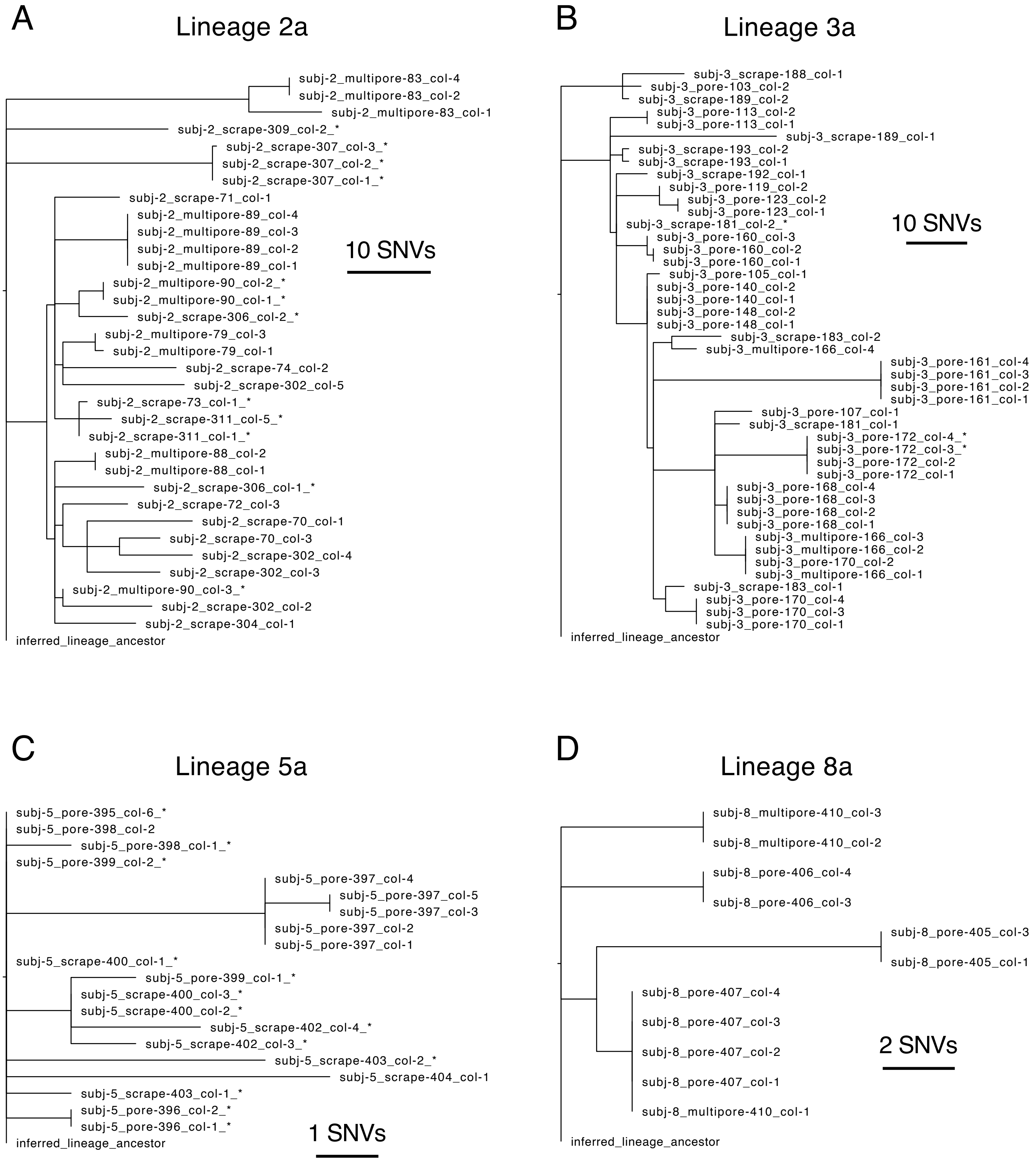


**Figure S6. Maximum parsimony trees of other lineages with pore samples from four different subjects.** These are the four largest lineages containing pore samples from four distinct subjects other than Subject 1. All lineages show patterns of low diversity within pores and pore-specific genotypes. Tip labels indicate individual colonies, named by their subject and sample of origin; asterisks in the colony name indicate the presence of a plasmid (see Figure S7).

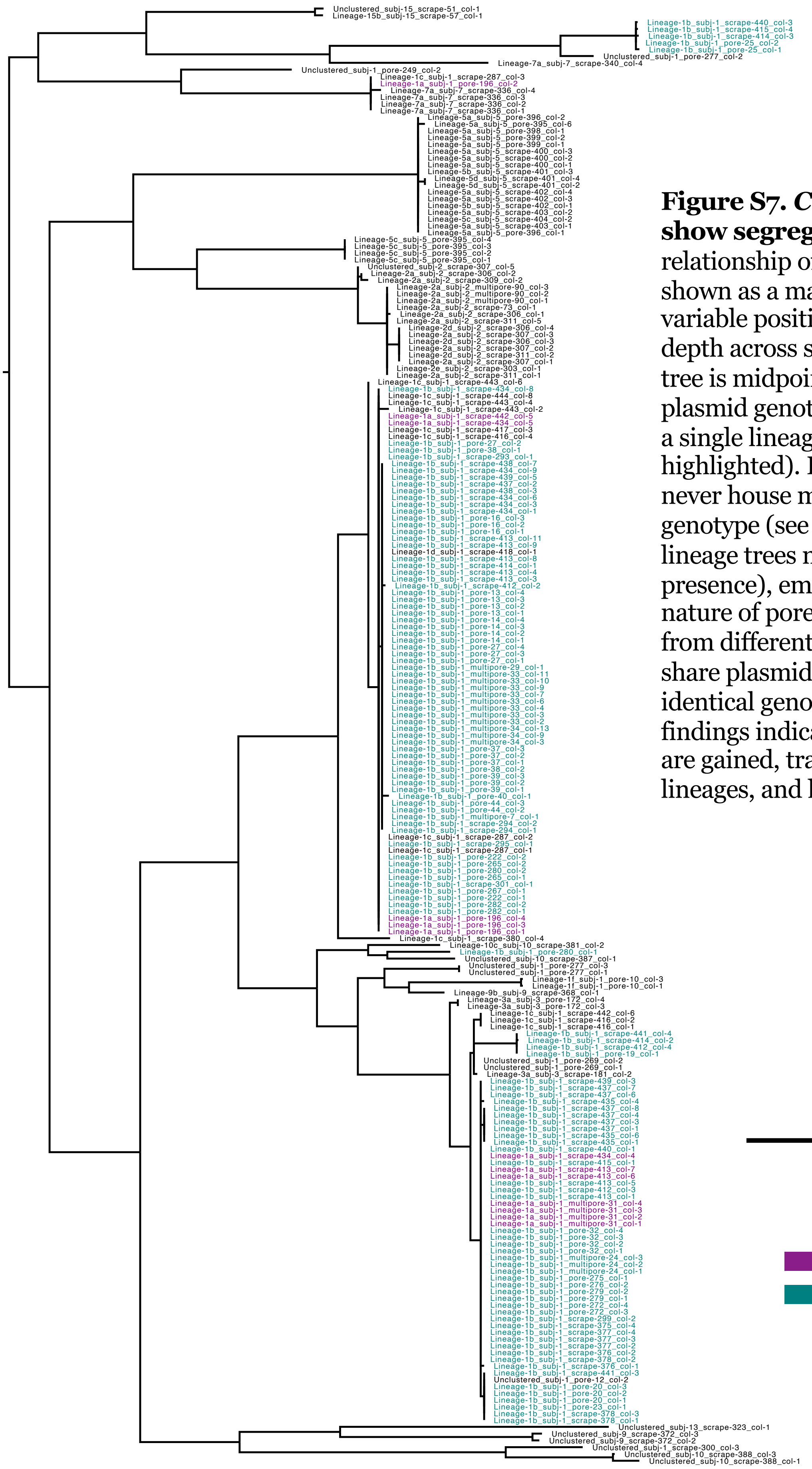

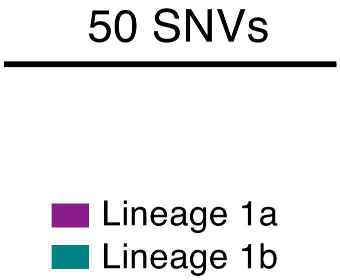


**Figure S7. *C. acnes* plasmids also show segregation by pore**. The relationship of plasmid genotypes is shown as a maximum parsimony tree of variable positions with sufficient read depth across samples (Methods). The tree is midpoint-rooted. Distant plasmid genotypes can be found within a single lineage (Lineages 1a and 1b highlighted). However, single pores never house more than one plasmid genotype (see also Figures S4-6 for lineage trees marked with plasmid presence), emphasizing the island-like nature of pores. In addition, colonies from different lineages on a person share plasmids with identical or near-identical genotypes. Together, these findings indicate that new plasmid types are gained, transferred between lineages, and lost on individual hosts.

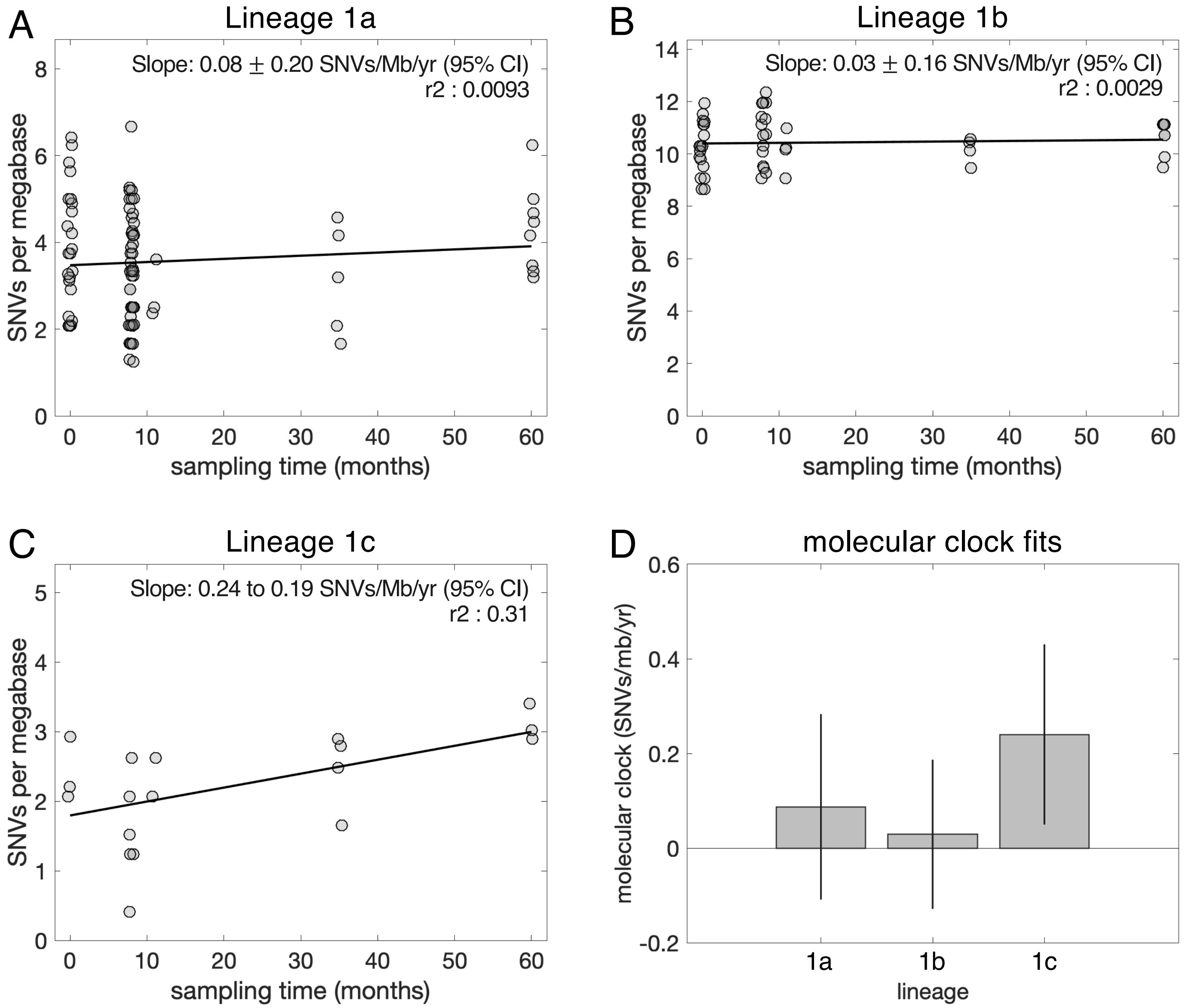


**Figure S8. A molecular clock for *C. acnes* evolution is not detectable over a 5-year time course.** We looked for a molecular clock signal in each of the three most abundant lineages on Subject 1, who was sampled longitudinally over five years. For each colony, we calculated the average number of mutations (SNVs) per 1 million base pairs (Mb) with sufficient coverage. We then averaged across colonies from a given sample, to account for the fact that colonies from the same sample are not independent. To assess the signal for mutation accumulation over time, we performed a linear regression between these sample averages and the time at which each sample was collected (note that some jitter is included in the time values displayed in the plots so that the point corresponding to each sample is visible). Fits for each lineage are shown in (A-C) and summarized in (D). We find no signal for a molecular clock within Lineages 1a and 1b, as indicated by a low $r^2$ value as well as by the 95% confidence interval of the slope including zero. This absence of a molecular clock signal suggests that the molecular clock for *C. acnes* on human skin is slower than could be captured by the length of the available time course or that either mutations/generation or generations/year are highly variable. For Lineage 1c, we find a detectable molecular clock signal of about 0.24 SNVs per megabase per year. While this value is consistent with values of molecular clocks from other species (Didelot et all, 2016), we opted not to convert numbers of mutations into amounts of time given the lack of signal in Lineages 1a and 1b.

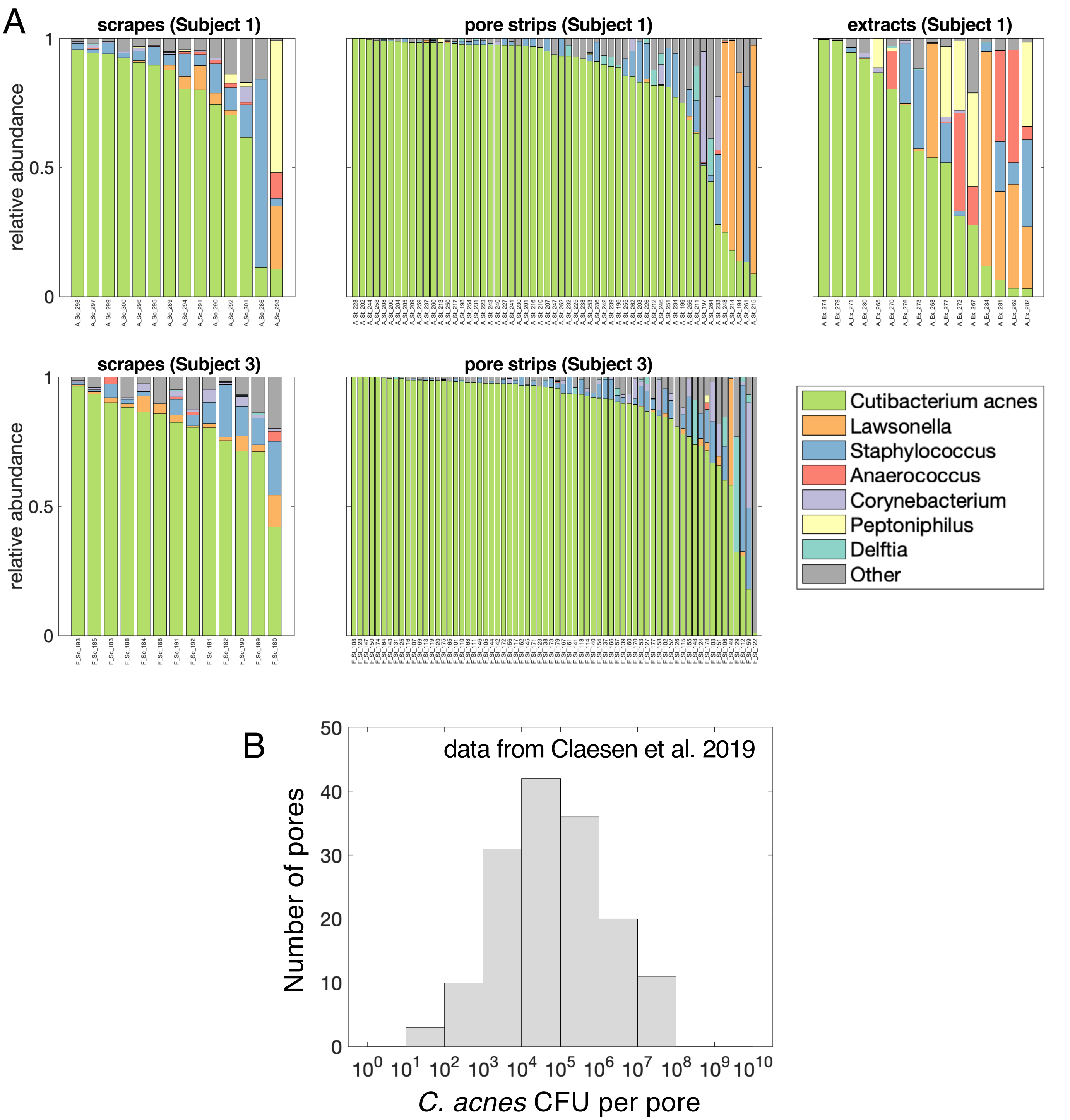


**Figure S9. *C. acnes* dominates within-pore bacterial populations.** (A) A subset of samples from two subjects was analyzed for community composition via 16S rRNA amplicon sequencing, in addition to colony-based *C. acnes* whole-genome sequencing (see Methods). Each species is colored in if its mean relative abundance across samples is >5% and each genus is colored if its max relative abundance across samples is >33%; everything else is categorized as "Other". Samples are segregated by collection method and subject. (B) Estimates of absolute abundances of *C. acnes* within individual pores derived from (Claesen et al., 2020). These estimates also agree with more limited data from our lab (data not shown).

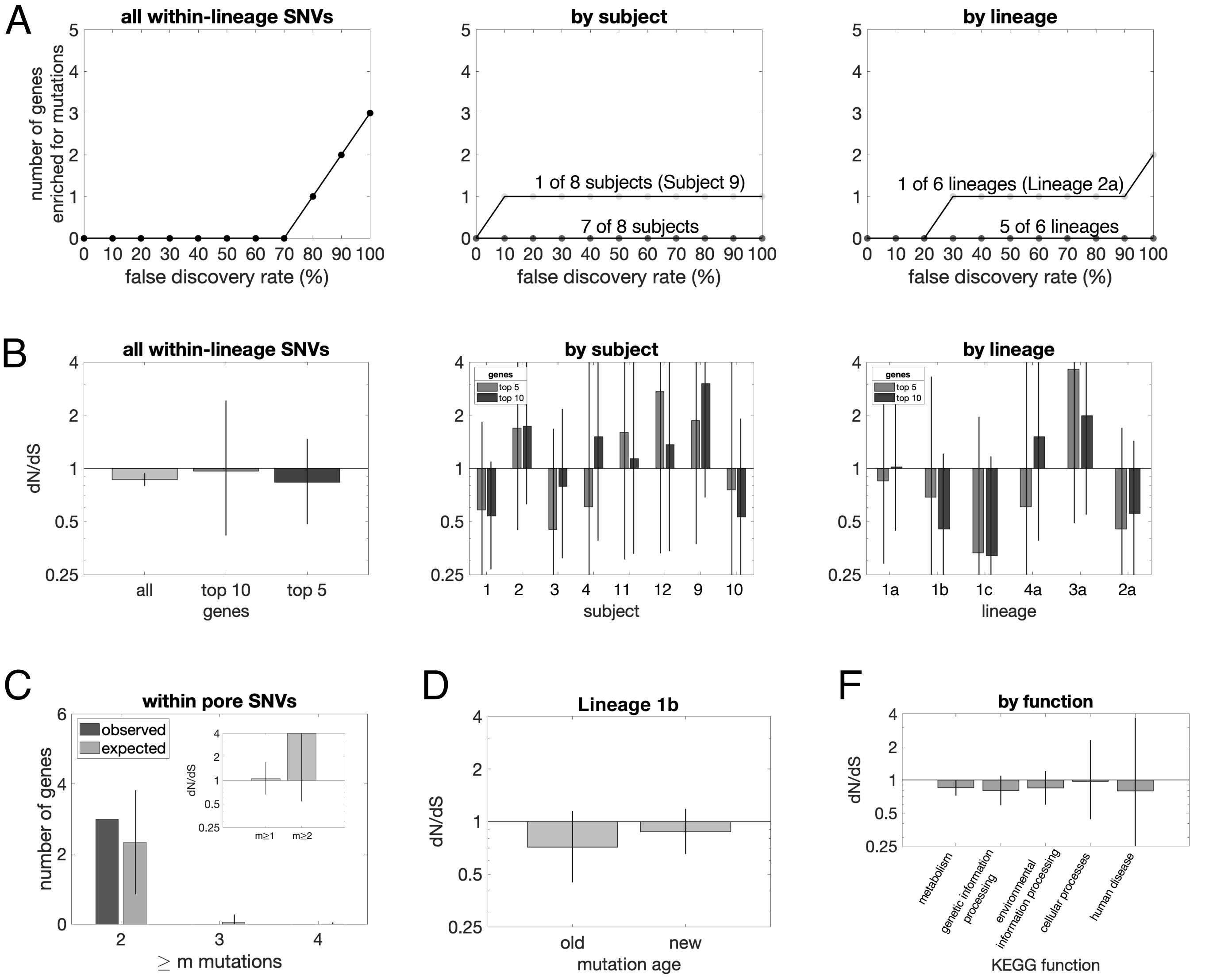


**Figure S10. No evidence of adaptive evolution among *de novo* SNPs.** (A) In order to search for parallel evolution, we calculated p-values for mutational enrichment for each gene (Methods) and performed the Benjamini-Hochberg procedure to correct for multiple hypothesis testing (treating each gene on the genome as a hypothesis). We plotted the number of genes detected for a range of FDRs for (i) the set of all observed *de novo* mutations, (ii) all subjects with at least 100 *de novo* mutations, and (iii) all lineages with at least 100 *de novo* mutations. We find two cases in which a gene is found with an FDR below 50%--an M18-family aminopeptidase on Subject 9 and a carbohydrate ABC transporter permease in Lineage 2a (each with 2 nonsynonymous mutations and 1 synonymous mutation, consistent with a neutral model). (B-E) We calculated dN/dS as in Figure 5B (Methods), but for different sets of mutations. (B) shows dN/dS for the top 10 and top 5 most mutated genes (according to Poisson p-values) across all mutations, by subject, and by lineage. (C) shows dN/dS for mutations inferred to have occurred inside pores (no parallel evolution detected with the Benjamini-Hochberg procedure; no enrichment of genes mutated multiple times relative to a neutral model; and no significant enrichment of nonsynonymous mutations on genes with multiple mutations, $p$=0.1); (D) shows dN/dS for older mutations on the long branches of Lineage 1b is not significantly different than newer mutations in the Lineage 1b subclades (Figure S5); and (E) shows dN/dS by KEGG function. Error bars represent 95% confidence intervals, demonstrating no significant signature for adaptive evolution.

A

$$\frac{\partial n}{\partial t} = D\frac{\partial^2 n}{\partial x^2} - v\frac{\partial n}{\partial x}$$

Drift-Diffusion equation where $n(x,t)$ is the probability that the cell is at pore depth $x$ at time $t$, with reflecting boundary conditions at the bottom of the pore ($x = 1000\mu m$) and absorbing boundary conditions at the top of the pore ($x = 0\mu m$)

B

| Parameter | Value | Source |
|---|---|---|
| Diffusion coefficient (*D*) of a cell in sebum | 650 μm²/day | Stokes-Einstein equation ($D = \frac{k_B T}{6\pi\eta r}$)<br>• Boltzmann constant $k_B$<br>• Body temperature $T = 37\text{C}$<br>• Sebum viscosity $\eta = 600$ millipoises {Butcher 1949}<br>• Cell radius $r = 0.5\ \mu\text{m}$ |
| Sebum flow speed (*v*) upward in the pore | -50 μm/day | Estimated from time for an extracted sebaceous follicle to refill (20 days) reported by (Plewig, 1974) for a pore of depth 1 mm |
| *C. acnes* doubling time ($t_g$) | 0.5 days | Estimated from max growth rates of *C. acnes* in laboratory aerobic conditions reported by (Cove et al.,1983) |

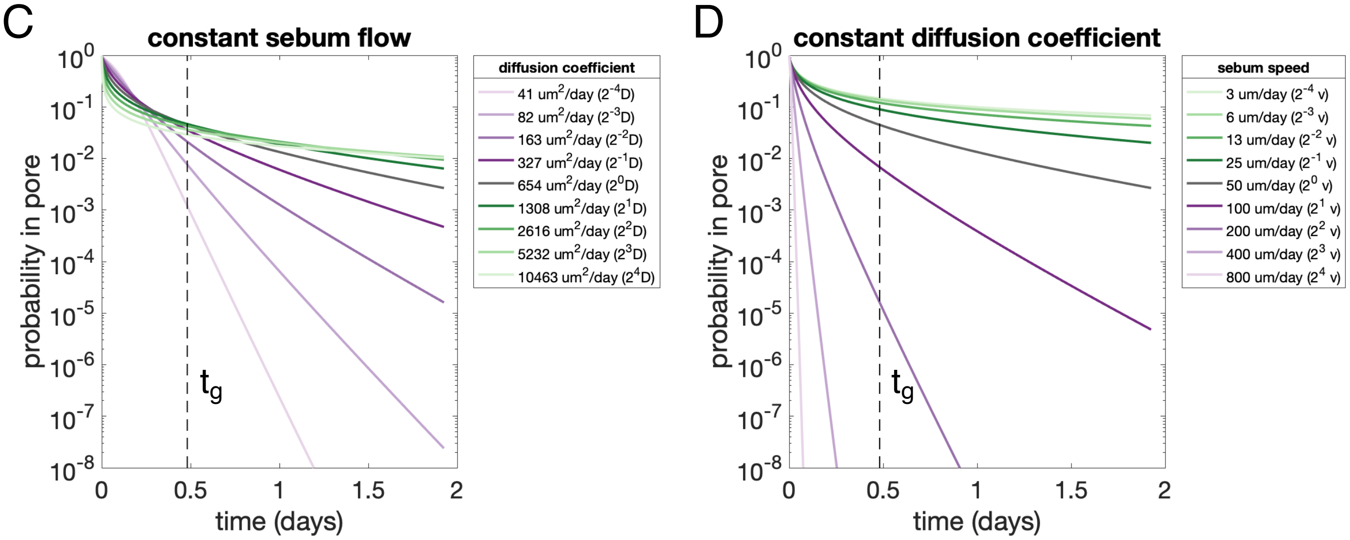


**Figure S11. Pore physiology may be responsible for neutral bottlenecking.** Individual cells attempting to colonize pores have a low likelihood of successfully colonizing a pore under reasonable assumptions. We modeled the diffusion of a cell through sebum as indicated in (A) and using conservative parameters estimated in (B), which are likely to overestimate the probability of a cell remaining in the pore, as a starting point. (C-D) We explored $2^4$-fold variation in either direction of the diffusion coefficient *D* and the sebum speed *v*, and we calculated the probability of a cell of a single cell remaining within the pore as a function of time. We assumed that a cell starts (at *t*=0) within in the top 10 um of the pore. (C-D) show how the probabilities change (C) for different diffusion coefficients and (D) for different rates of sebum flow. These probabilities are particular sensitive to sebum flow rate. While the number of cells attempting to colonize a pore at a given time is unknown, these low likelihoods of successful colonization provide many generations during which a single-cell colonizer can expand to dominate the pore before the next successful colonization event. In addition, a cell that reaches the lower depths of the pore will have an advantage over later colonizers because of the anaerobic environment. Moreover, diffusion will likely be substantially slower due in a partially or fully colonized pore, where cells that are already in the pore create obstacles to other cells--making it even more difficult for a new colonizer to penetrate deeper into the pore.

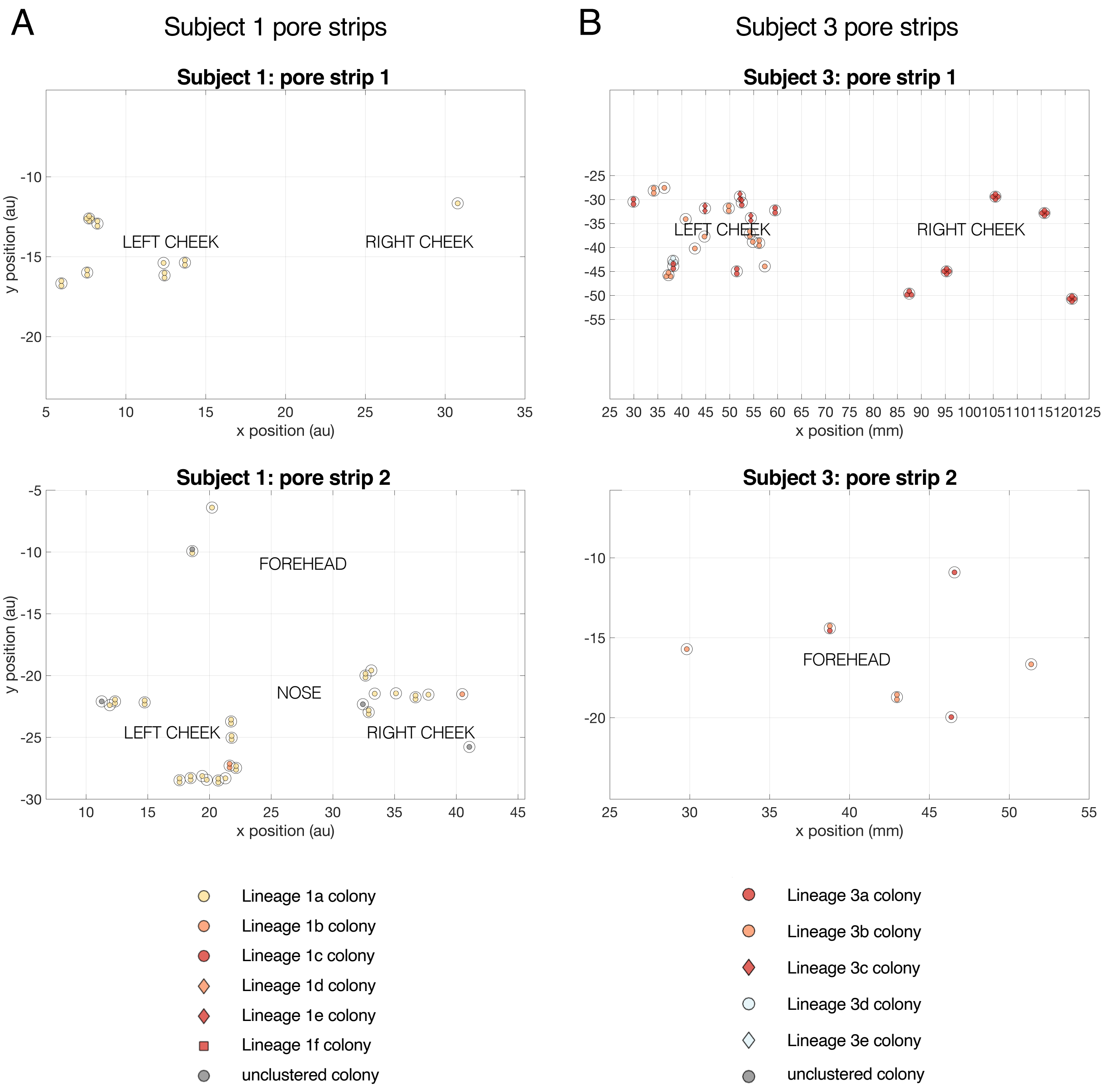


**Figure S12. Spatial coordinates and lineages for pore strip colonies**. Pore strip samples from Subject 1 (A) and Subject 3 (B) enabled us to collect colonies from pores with defined spatial coordinates. Each sample derived from a single follicle is denoted by an open circle. Each colony from that sample is represented by a symbol inside that circle, where the color represents the strain type (same color scheme as Figures 2, 3, and S3) and the shape represents the lineage. Approximate facial regions are marked with text labels. Lineages are ordered by abundance across all colonies from the subject (even if that lineage was not represented in pore strip colonies).

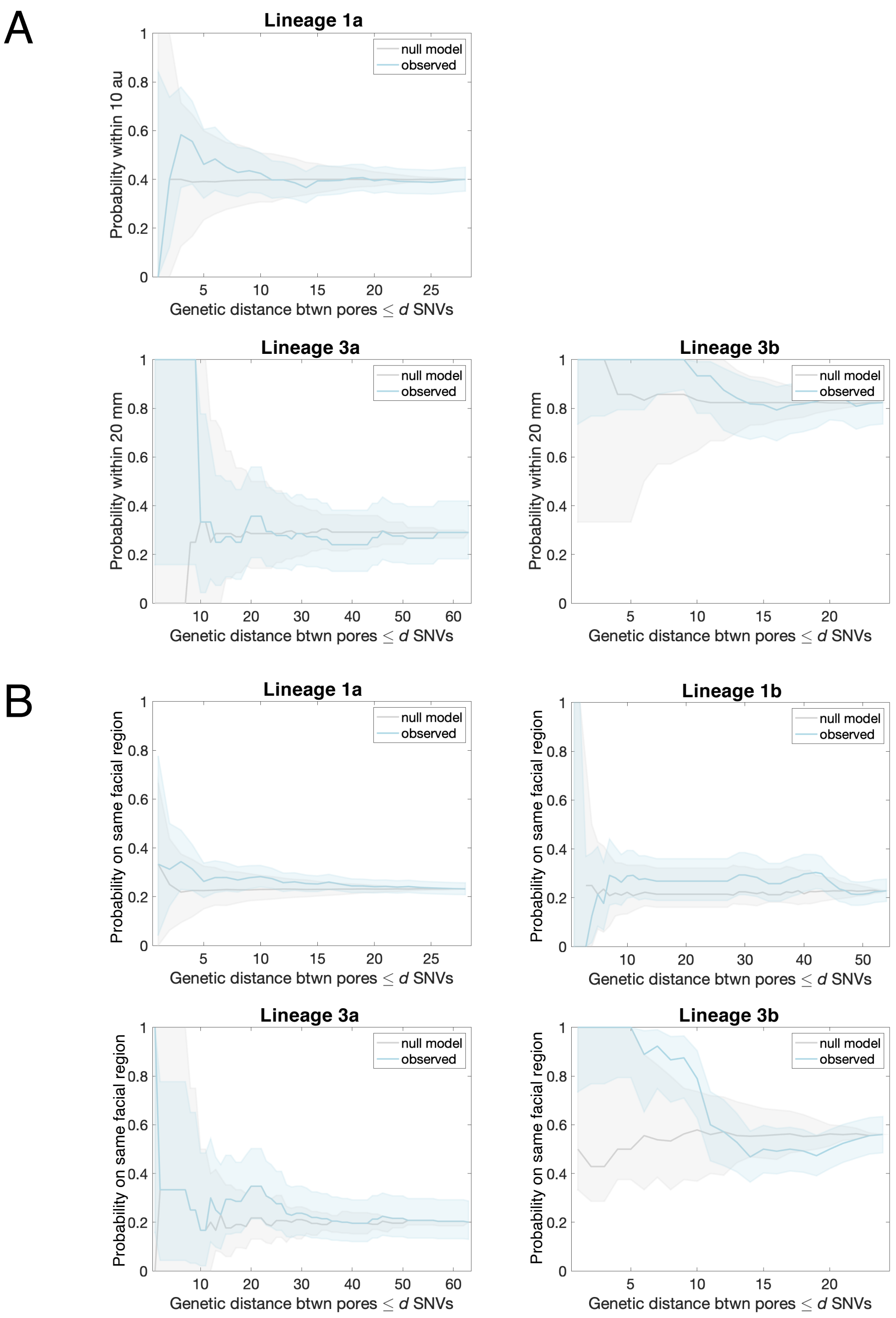


**Figure S13. Lineages spread across the face faster than they accumulate mutations**. (A) In order to determine if physical distance between pores was related to genetic distance between pores, we computed the the probability that pairs of pores harboring colonies from the same lineage within *d* SNVs of each other were co-localized (within 20 mm for Subject 3; within 10 arbitrary units for Subject 1, which is approximately equivalent to 20 mm). We compared this probability to that expected from a null model, where the spatial coordinates of pores were re-shuffled. Plots show 95% CIs (binomial distribution). Only pore sample originating from a single follicle and whose colonies were monophyletic were used in this analysis, so that we could compute the genetic distance between pores as the number of SNVs between inferred pore ancestors (Methods); lineages with at least 10 pores with spatial coordinates are included. (B) We repeated this analysis including colonies from pore extracts subject to the same criteria as noted above, computing the probability that genetically related pairs of pores were confined to the same facial skin region (see Figure 3). In both cases, we did not detect spatial confinement, suggesting that lineages spread across the face faster than they accumulate mutations.

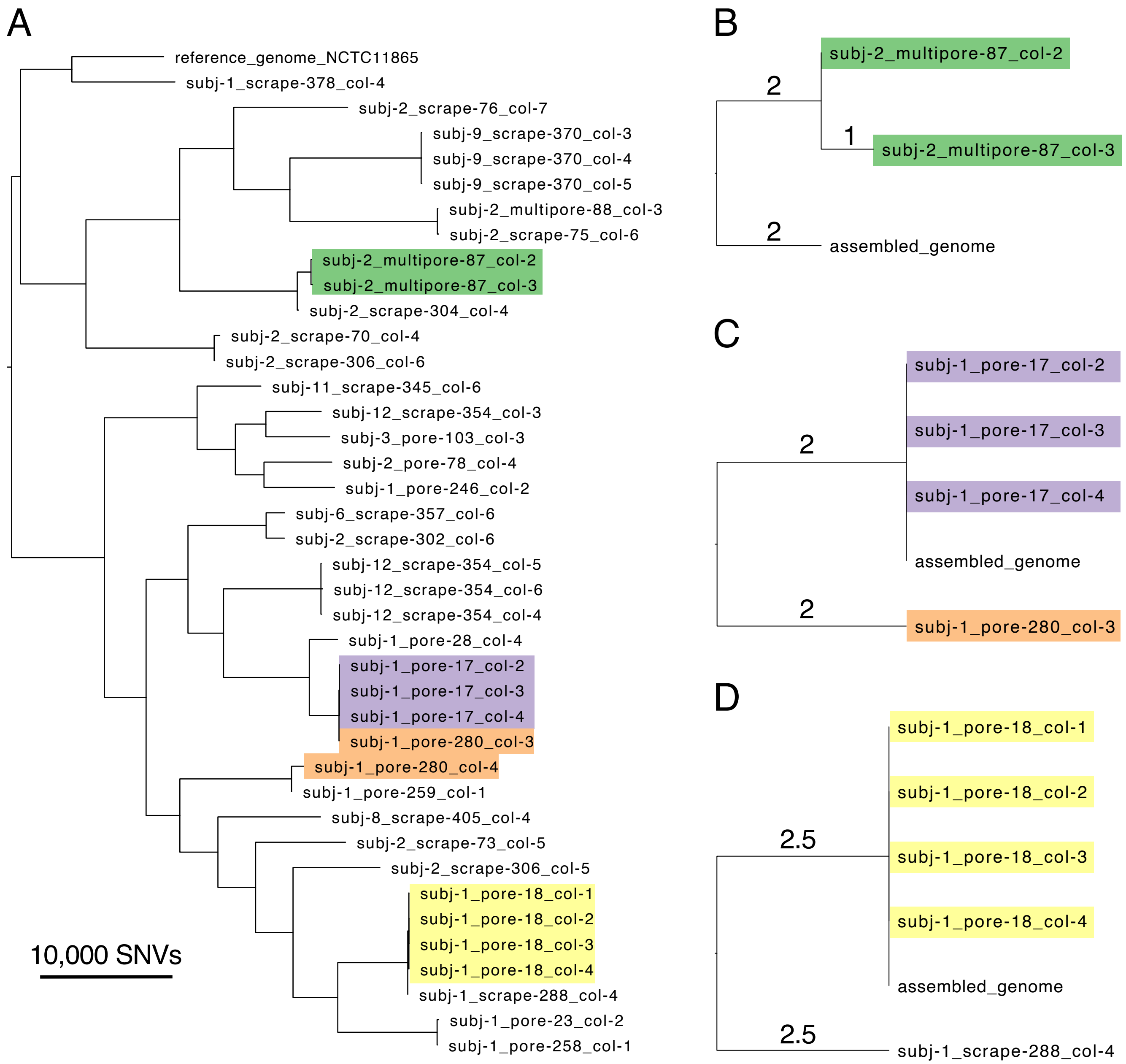


**Figure S14. Limited genetic diversity among *Cutibacterium granulosum* colonies originating from the same pore.** While aiming to sequence *C. acnes* colonies, we sometimes cultured colonies of other species, and 50 colonies were identified as *C. granulosum*. (A) SNVs were analyzed using an alignment-based approach to reference genome NCTC11865 and evolutionary reconstruction was performed, using a similar approach as for *C. acnes* (Methods). Maximum parsimony of the 39 *C. granulosum* colonies that passed quality filtering shows the presence of distinct lineages on each person. Note that genetic distances are approximate, as no effort was made to distinguish variable gene content or recombination from SNVs. Cases where multiple colonies originate from the same pore sample are highlighted. (B-D) In order to investigate the within-pore diversity at a finer scale, we assembled genomes for each of the three cases where colonies from the same pore were monophyletic and repeated evolutionary reconstruction (Methods). Branch lengths are annotated in SNVs. These trees show that *C. granulosum* colonies from the same pore share nearly identical genomes. (A-D) All trees are midpoint rooted.
